# Impact of high fat diet and exercise on bone and bile acid metabolism in rats

**DOI:** 10.1101/2024.01.07.573992

**Authors:** Nerea Alonso, Gunter Almer, Maria Donatella Semeraro, Giovanny Rodriguez-Blanco, Günter Fauler, Ines Anders, Gerald Ritter, Annika vom Scheidt, Niels Hammer, Hans Jürgen Gruber, Markus Herrmann

## Abstract

**Background:** Bile acids help facilitate the intestinal lipids absorption, and have endocrine activity in glucose, lipid and bone metabolism. Obesity and exercise influence bile acid metabolism and have opposite effects in bone. This given study investigates if regular exercise helps mitigate the adverse effects of obesity on bone, potentially by reversing alterations in bile acid metabolism.

**Methods:** Four-month-old female Sprague Dawley rats either received high fat diet (HFD) or chow-based standard diet (lean controls). During the 10-month study period, half of the animals performed 30 min of running at moderate speed on five consecutive days followed by two days of rest. The other half was kept inactive (inactive controls). At study end, bone quality was assessed by micro computed tomography and biomechanical testing. Bile acids were measured in serum and stool.

**Results:** HFD feeding was related to reduced trabecular (−33%, p=1.14×10^-7^) and cortical (−21%, p=2.9×10^-8^) bone mass, and lowered femoral stiffness (12-41%, p=0.005). Furthermore, HFD decreased total bile acids in serum (−37%, p=1.0×10^-6^), but increased bile acids in stool (+2-fold, p=7.3×10^-9^). These quantitative effects were accompanied by changes in the relative abundance of individual bile acids. The concentration of serum bile acids correlated positively with all cortical bone parameters (r=0.593 - 0.708), whilst stool levels showed inverse correlations at cortical (r=-0.651 - −0.805) and trabecular level (r=-0.656 - −0.750). Exercise improved some trabecular and cortical bone quality parameters (+11-31%, p=0.043 to 0.001) in lean controls, but failed to revert the bone loss related to HFD. Similarly, changes in bile acid metabolism were not mitigated by exercise.

**Conclusion:** Prolonged HFD consumption induced quantitative and qualitative alterations in bile acid metabolism, accompanied by bone loss. Tight correlations between bile acids and structural indices of bone quality support further functional analyses on the potential role of bile acids in bone metabolism. Regular moderate exercise improved trabecular and cortical bone quality in lean controls but failed mitigating the effects related to HFD in bone and bile acid metabolism.

## 1. Introduction

In developed countries, large parts of the population adhere to a hypercaloric diet that contains an excessive amount of animal-derived dietary fat. Especially when combined with physical inactivity, a high nutritional fat content is associated with increased body weight, visceral fat accumulation and metabolic dysfunction (1, 2). Obesity, which affects approximately 30% of the world’s population (3), promotes several chronic diseases, including cardiovascular disease, hypertension, diabetes and liver disease that increase morbidity and mortality (4). Osteoporosis, another common condition in aging Western societies, has also been linked to obesity, but existing evidence is inconsistent. Early studies suggested higher bone mineral density (BMD) in obese individuals as result of increased mechanical loading (5–7), whereas later studies in obese individuals reported comparable or reduced bone quality with increase porosity compared to controls (8–12).

A high nutritional fat intake increases the secretion of bile acids into the gut, where they function as emulsifiers that facilitate the absorption of lipids and fat-soluble vitamins (13, 14). Today it is well established that bile acids also possess endocrine activity that modulates glucose and lipid homeostasis (15). Furthermore, their possible contribution to different metabolic processes has attracted interest in late years as a regulatory pathway in some obesity-associated comorbidities, such as type 2 diabetes or chronic liver disease (16). Previous studies also suggest a regulatory role of bile aids in bone metabolism. In postmenopausal women, low serum bile acid concentrations are associated with osteoporosis (17). Furthermore, the serum concentration of total bile acids is positively related to BMD (18), and in osteoporotic patients, composition of the serum bile acid pool was found to be substantially altered (19). The endocrine actions of bile acids are meditated by several receptors like the Farnesoid X receptor (FXR) and the Takeda G protein-couple receptor 5 (TGR5), which differ in their affinity for individual bile acid species (20). The importance of both receptors for health has been substantiated in knockout mice, which are characterized by bone loss and increased osteoclastic bone resorption (21–23).

Bile acids are catabolic products of cholesterol that are formed in the liver and subsequently secreted into the bile duct. Synthesis and metabolism of bile acids are tightly regulated in the liver and the gut in order to ensure appropriate digestion without cytotoxic effects. Bile juice contains 97-98% water and less than 1% bile acids. After a meal, bile juice is secreted into the gut where bile acids help to absorb lipids and fat-soluble vitamins (13, 14). Hepatocytes synthesize the primary bile acids cholic acid (CA) and chenodeoxycholic acid (CDCA), which are subsequently conjugated with taurine and glycine. In the gut, bile acids are further metabolized by intestinal microbiota forming secondary bile acids, like lithocholic (LCA), deoxycholic (DCA), ursodeoxycholic (UDCA) and hyodeoxacholic acid (HDCA). In mice, CDCA is also used as a substrate to generate murine-specific bile acids named muricholic acids (MUA) (14, 24, 25). Like primary bile acids, also secondary bile acids are conjugated with taurine or glycine. Normally, 95% of the bile acids return to the liver via the enterohepatic circulation so that only small quantities are lost through the stool, and thus have to be replaced by de novo synthesis (26, 27). However, metabolic conditions like obesity can substantially impact bile acid homeostasis including alterations in composition of the bile acid pool. For example, high dietary fat consumption enhances hepatic synthesis and secretion of bile acids with higher intestinal amounts that are metabolized by gut microbiota to secondary species (28). In osteoporotic patients, primary bile acid synthesis seems to be shifted from the classic pathway to the alternative one (19). The classical pathway generates primarily CA from cholesterol, whereas the alternative pathway results in the production CDCA (29). Such changes may be important for bone health as previous studies suggest differential effects of individual bile acid species in bone, sometimes with opposite directions (30, 31).

Regular exercise, an established way to reduce body weight and the risk of obesity-associated diseases (32), has beneficial effects on bone health that aid the prevention and treatment of osteoporosis (33, 34). In people over 65 years of age, regular exercise is associated with higher BMD at hip and lumbar spine, with the more extensive programs that include a mix of exercises being most effective (35). Limited evidence also suggests regulatory effects of physical activity on bile acid metabolism. A lower serum concentration of total bile acids has been shown in amateur runners after a half-marathon run (36) whilst resistance and endurance exercise showed more divergent effect on circulating concentrations of specific bile acids in moderately trained males (37). The effects of exercise on bile acid metabolism seem to be of quantitative and qualitative nature, with a different composition of the circulating bile acid pool in physically active individuals (37, 38). LCA, for example has been shown to be higher in exercising individuals.

Considering the opposite effects of obesity and exercise in bone, we hypothesised that regular exercise could mitigate the adverse effects of obesity on bone, potentially by reversing alterations in bile acid metabolism. The present study aimed to address this question in a rat model where animals were fed for an extended period of time with a hypercaloric high fat diet (HFD) or a chow-based standard diet. To explore potential exercise effects, half of the animals performed regular moderate treadmill running, whereas the other half was kept sedentary. Following the intervention, bone structure and biomechanical properties were analysed together with an extensive panel of bile acids in serum and stool.

## 2. Methods

### 2.1. Animals

Four-month-old Sprague Dawley female rats were purchased from Janvier Labs (Le Genest-Saint-Isle, France) (n=96) and kept at the Biomedical Research, Medical University of Graz (Graz, Austria) as previously described (39). In summary, animals were maintained in groups of three per cage on constant housing conditions (12h light/12h dark cycles). The average weight of the animals was 300 g at baseline. This strain is naturally prone to develop benign cyst tumours and any animal presenting a tumour was excluded from the analysis. All animal experiments were performed following permission from the Austrian Federal Ministry of Education, Science and Research (GZ: 66.010/0070-V/3b/2018). They complied with the ARRIVE guidelines (https://www.nc3rs.org.uk/arrive-guidelines).

### 2.2. Diet and exercise

Ninety-six animals were randomly split in two groups (n=48 per group) and were fed for 10 months either with a standard diet or a high fat diet (HFD). The standard diet (Altromin, Lage, Germany) contained 11% fat and providing 3,226 kcal/kg, whereas the HFD was based on purified beef-tallow with 60% fat (8.3% of C16:0, 6.1% of C18:0 and 12.3% of C18:1), which provided 5,150 kcal/kg (ssniff, Soest, Germany). Food and water were provided ad libitum. Both diet groups were subdivided randomly into an inactive control group and an exercise group (n=24 animals per subgroup). The inactive control groups did not perform any kind of exercise or regular physical activity. In contrast, animals in the two exercise groups underwent a moderate standardised exercise program consisting in 30-min running on a treadmill (Panlab, Barcelona, Spain) at constant speed of 30 cm/s on five consecutive days per week followed by two days of rest. All animals of the two exercise groups completed their running protocol within a window period of 4 h starting at 10:00am. At the end of the study period, animals were sacrificed by heart puncture under deep isofluorane anesthesa (Forane, Abbott, Austria).

### 2.3. Micro computed tomography (microCT)

Lower limbs of animals from the four experimental groups (n=10 per group) were dissected, soft tissue was removed and they were fixed in 4% paraformaldehyde in PBS. Tibial cortical and trabecular bone was scanned *ex vivo* using a SkyScan 1276 (Bruker, Kontich, Belgium) microCT device at a resolution of 10 µm (70 kV, 200 μA, 0.5-mm aluminium filter, rotation step size 0.25 degrees) at the Division of Biomedical Research, Medical University of Graz (Graz, Austria). Image reconstruction was performed using the Bruker Skyscan NRecon v1.7.4.2 software with a beam hardening of 30%. Trabecular measurement was performed at 100 slices (1 mm) from the growth plate and then 100 slices (10 µm each) were selected for analysis. Cortical bone was analysed at 12.5 mm from the growth plate where six regions of 100 slices each with a separation of 35 slices were selected using the Skyscan CTAn software. All bone parameters were adjusted by body weight.

### 2.4. Biomechanical analysis

To assess the influence of the intervention on whole bone biomechanics, femora from five rats per group were subjected to three-point loading. The distance between bearing regions was adapted to equal half the length of each femur, in accordance with recommendations from Prodinger et al (40). Quasi-static three-point bending tests were performed on a uniaxial mechanical testing machine (Z020, ZwickRoell GmbH & Co. KG, Ulm Germany) equipped with a 2.5-kN load cell (41). Samples were subjected to a pre-load of 2 N following a waiting time of 5 s, each sample was loaded with 0.1 mm/s until fracture. Force and deflection were recorded continuously throughout the trial. Main outcome parameters included maximum flexural stress and Young’s modulus. For maximum flexural stress calculation, the inner and outer radius of the femora were determined via microCT and the femoral shaft was approximated as a hollow cylinder. Young’s modulus was calculated in the elastic quasi-linear region of the loading curve.

### 2.5. Extraction of bile acids

Blood was collected by heart puncture and serum was obtained by centrifugation of the samples at 2000g for 12 min at room temperature. Stool samples were collected from the descending colon at sacrification and, both serum and stool were kept at −80 °C for analysis. Animals who developed tumours during the intervention period were excluded from analysis. The exact number of remaining animals in each group is indicated in each analysis. Bile acids were extracted as previously described (42). In summary, 10 µL (0.2 nmol each) internal standards purchased from Sigma Aldrich (Taufkirchen, Germany), Steraloids (Newport, RI, USA) or synthesised in house were added to the serum and spin-vortexed. Deproteination was performed by adding 400 µL of acetonitrile, followed by vortex and centrifugation at 3200 g for 12 min at room temperature. The supernatant was dried at 50 °C under constant flow of nitrogen. Subsequently, samples were reconstituted and transferred to autosampler vials.

Bile acids from stool samples were extracted following previous publications (42, 43). Snap frozen stool samples (10 mg) were incubated with NaOH (0.1 M, 2 ml) for 60 min at 60 °C. After adding 4 ml of aqua dest, samples were homogenised and protein was denatured by adding 80% v/v of acetonitrile for 20 min at room temperature. Internal standards were added as indicated above. Proteins were removed by 20 min of centrifugation at 20,000 g. The supernatant was dried at 50 °C under nitrogen flow, and the resulting pellet was re-dissolved in 4 ml of ammonium acetate and purified using C18 reversed phase SPE cartridges. 20 ml of aqua dest were used to remove hydrophilic material, whereas lipophilic component was removed with 10 ml of hexane. Finally, bile acids were eluted with 2 ml of methanol.

### 2.6. Mass spectrometry analyses

Twenty-nine bile acids, including primary, secondary, and conjugated species (Suppl Table S1) were analysed by liquid chromatography-high resolution-mass spectrometry (LC-HR-MS). Chromatography of 10 µL of each sample was performed using a Nucleoshell C18 reversed phase column (Macherey-Nagel, Düren, Germany) for human bile acids. Murine-specific bile acids were analysed using a Kinetex pentafluorophenyl (PFP) column. Separation was performed using aqua dest with 1.2% v/v formic acid and 0.38% w/v ammonium acetate, and elution was carried out using methanol with 1.3% v/v formic acid and 0.38% ammonium acetate. Gradient settings are described elsewhere (42). Analysis was performed on a Q Exactive hybrid quadrupole-orbitrap mass spectrometer (Thermo Fisher Scientific, Massachusetts, US) with an ESI ion source in negative ionisation mode. Instrument settings were described in detail previously (42). The limit of quantitation of the mass spectrometer was 0.001 µmol /L for all bile acid species. Any value below this threshold was not quantitated and thus excluded from statistical analyses.

### 2.7. Statistics

Parametric distribution of each parameter was tested using Shapiro-Wilk normality test in IBM SPSS v27. Changes in bone parameters or bile acid levels between groups were assessed by two-way ANOVA. In case of a non-parametric distribution, the ANOVA test was performed on previously calculated ranks. Comparisons between two groups were performed by Student’s t-test for normally distributed parameter or Mann-Whitney test for not normally distributed ones. Correlation analyses were performed by Person’s correlation for parametric variables and Spearman’s correlation for non-parametric variables. *P* values below the threshold for Bonferroni correction for multiple testing were considered significant. Specific thresholds are mentioned in each statistical analysis. Graphs were generated using GraphPad Prism v9.2.0 (California, US).

## 3. Results

### 3.1. Exercise cannot revert long-term-HFD-related bone loss

At the end of the 10-month intervention period, microCT analyses (n=10/group) revealed a significant deterioration of bone architecture in HFD animals (Figure 1 and Suppl. Figures S1-2). When compared to control animals, trabecular bone mass (BV/TV) was reduced by 33%, trabecular surface density (BS/TV) by 29%, trabecular thickness (Tb.Th) by 24% and trabecular number (Tb.N) by 29% (Figures 1 and Suppl. Figure S1). In contrast, regular running exercise was related to increased trabecular parameters when compared with sedentary controls on standard diet (BV/TV + 31%, p=0.021; BS/TV +16%, p=0.025; Tb.Th +19%, p=0.043; Tb.N+20%, p=0.016). However, exercise could not revert HFD-related bone loss at trabecular level (Figures 1 and Suppl. Figure S1). microCT analyses of cortical bone showed a loss in HFD animals when compared to sedentary control animals on standard diet. HFD reduced cortical thickness (Ct.Th) by 25%, cortical area (Ct.Ar) by 24%, cortical area fraction (Ct.Ar/Tt.Ar) by 21%, periosteal perimeter (Ps.Pm) by 17%, endocortical perimeter (Ec.Pm) by 15%; maximal inertia (I_max_) by 19%, minimal inertia (I_min_) by 24% and polar moment of inertia (J) by 20% (Figures 1 and Suppl. Figure S2). Exercise improved some cortical parameters when compared to controls on standard diet (Ct.Ar +13%, p=0.003; Ps.Pm +11%, p=0.001; Ec.Pm +15%, p=0.002; I_max_ +23%, p=0.005; I_min_ +18%, p=0.003; J +22%, p= 0.003). Similar to trabecular bone, exercise did not revert cortical bone loss in HFD fed animals (Figures 1 and Suppl Figure S2).

**Figure 1.**
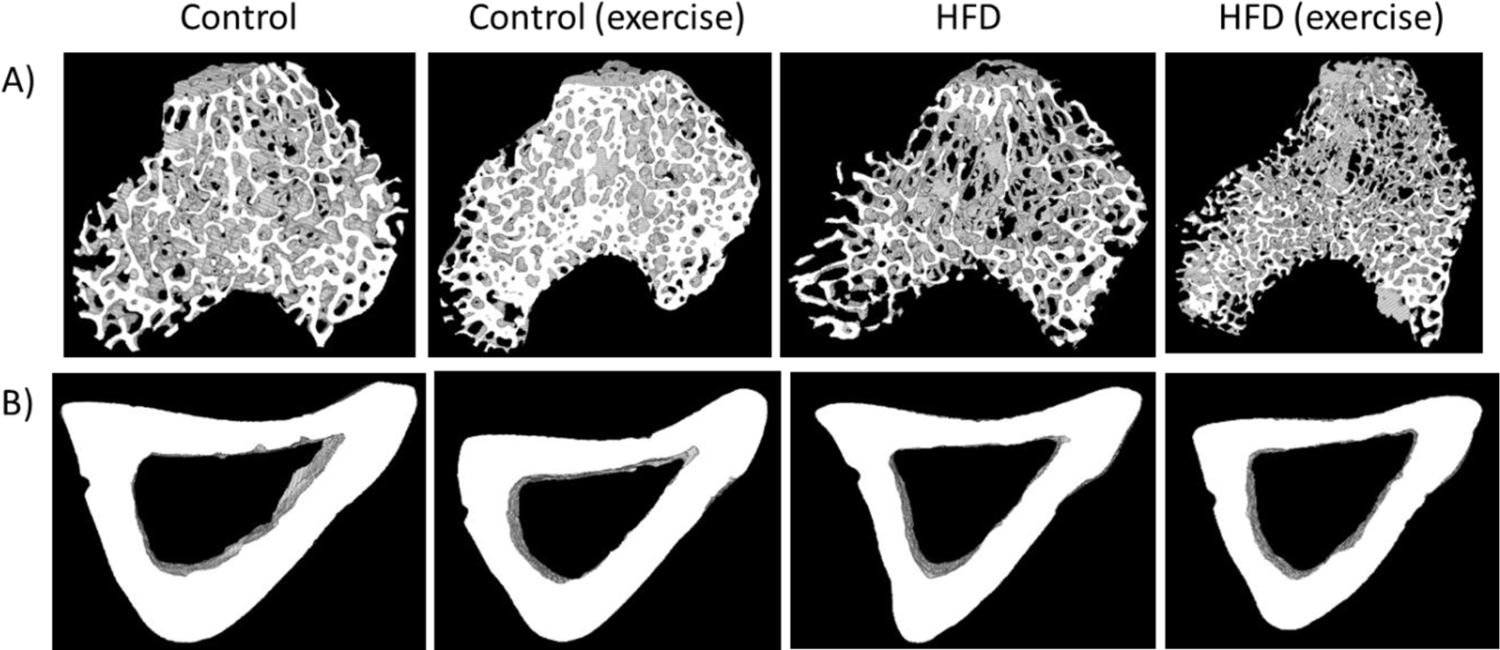
Exemplary microCT analysis of A) trabecular and B) cortical tibia of the animals from the four intervention groups.

### 3.2. Long term HFD and exercise have effect on bone mechanical properties

The mechanical properties of femurs from five animals per subgroup were analysed by 3-point bending test. Mean maximal force and flexural stress as well as Young’s modulus were calculated for each of the four groups. Results showed that both, HFD and exercise altered the mechanical properties of the bone. Both interventions decreased the maximal force that was needed to fracture the bone. This effect was particularly pronounced in exercising HFD animals, with a reduction of 41%. However, the interaction between HFD and exercise did not show any significant effect. The maximal flexural stress was 40% lower in exercising HFD animals compared to controls. No changes were observed for Young’s modulus (Figure 2).

**Figure 2.**
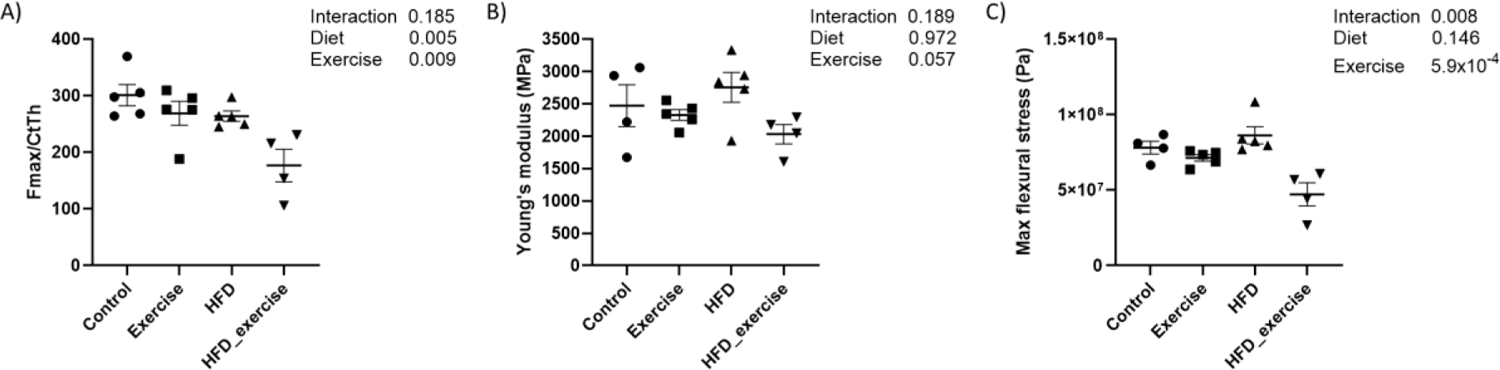
Biomechanical properties of rats treated with high fat diet and subject to exercise compared with controls: A) maximal force corrected by cortical thickness; B) Young’s modulus; C) maximal flexural stress.

### 3.3. HFD-related alterations of serum bile acid profile cannot be reverted by exercise

HFD animals showed an altered composition of the serum bile acid pool (Suppl. Table S1) when compared to controls on standard diet (Figure 3). The concentration of total bile acids in HFD animals was 37% lower than in controls. The greatest reductions were seen for free (−75%), primary (−68%) and 12-α-hydroxylated (12-α-OH) (−58%) bile acids (Table 1). Overall, total conjugated bile acids were reduced by 56% (p=0.002) in HFD animals. However, analysis of the individual conjugated bile acid species showed a variable response to HFD with reductions of glycocholic acid (−91%), taurochenodeoxycholic acid (−67%), glycoursodeoxycholic acid (−60%) and murine tauro alpha muricholic acid (−60%), but an increase of 4.5-fold of taurohyodeoxycholic acid. The total secondary bile acid concentration did not change in response to HFD, despite significant alterations of individual species, such as ursodeoxycholic acid (−85%) and lithocholic (−82.4%). The complete bile acid profile of all groups is shown in Table 1.

**Figure 3.**
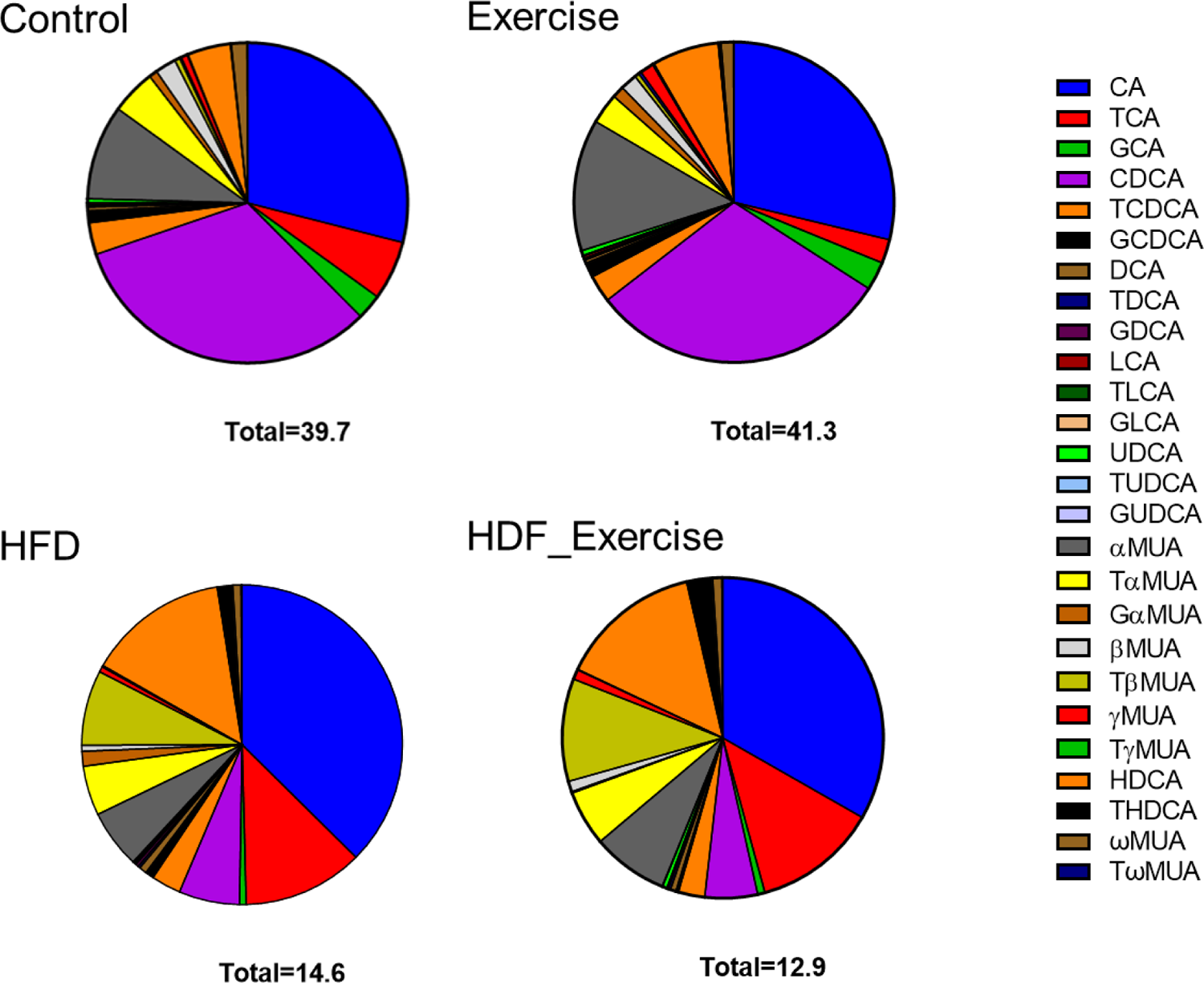
Distribution of bile acids (median) in the serum of each of the intervention groups and controls. Total concentration of all species of bile acids is shown at the bottom of each graph.

**Table 1.** Significant changes in bile acids (µM) in serum associated with high fat diet and physical activity (threshold for significance p =7.0×10^-4^)

| Bile acid | Control |  | Exercise |  | HFD |  | HFD & Exercise |  | Two-way ANOVA p |  |  |
| --- | --- | --- | --- | --- | --- | --- | --- | --- | --- | --- | --- |
|  | N | Median [IQR] | N | Median [IQR] | N | Median [IQR] | N | Median [IQR] | HFD | Exercise | Interaction |
| **Total** | 21 | 39.742<br>[19.451 - 72.835] | 23 | 41.322<br>[18.440 - 79.039] | 16 | 14.570<br>[9.514 - 20.640] | 12 | 12.858<br>[4.721 - 18.966] | $1.0 \times 10^{-6}$ | 0.600 | 0.665 |
| **Conjugated** | 21 | 8.242<br>[5.133 - 12.355] | 23 | 5.905<br>[3.422 - 8.506] | 16 | 3.671<br>[1.839 - 7.333] | 12 | 3.132<br>[1.433 - 6.188] | 0.002 | 0.237 | 0.536 |
| **Free** | 21 | 33.621<br>[14.310 - 58.860] | 23 | 36.497<br>[13.479 - 51.702] | 16 | 8.357<br>[3.469 - 14.666] | 12 | 4.982<br>[0.733 - 12.151] | $2.4 \times 10^{-7}$ | 0.600 | 0.452 |
| **Primary** | 21 | 35.461<br>[18.838 - 61.398] | 23 | 35.156<br>[17.508 - 66.429] | 16 | 11.360<br>[7.431 - 16.941] | 12 | 9.053<br>[3.185 - 15.714] | $6.4 \times 10^{-8}$ | 0.437 | 0.629 |
| **Secondary** | 21 | 3.082<br>[0.761 - 8.755] | 23 | 3.662<br>[0.945 - 8.885] | 16 | 2.776<br>[1.485 - 3.827] | 12 | 2.500<br>[1.517 - 3.756] | 0.278 | 0.741 | 0.573 |
| **12-α-OH** | 21 | 16.230<br>[9.420 - 27.019] | 23 | 12.237<br>[8.287 - 20.559] | 16 | 7.013<br>[5.231 - 12.021] | 12 | 5.001<br>[1.887 - 11.426] | $9.6 \times 10^{-5}$ | 0.189 | 0.884 |
| **CA** | 21 | 11.397<br>[6.426 - 21.177] | 23 | 10.090<br>[5.583 - 15.015] | 15 | 5.272<br>[2.707 - 7.380] | 9 | 4.124<br>[0.630 - 7.124] | $2.0 \times 10^{-5}$ | 0.438 | 0.861 |
| **GCA** | 21 | 1.009<br>[0.384 - 2.151] | 23 | 1.009<br>[0.689 - 1.507] | 16 | 0.093<br>[0.059 - 0.254] | 11 | 0.091<br>[0.073 - 0.207] | $4.3 \times 10^{-12}$ | 0.804 | 0.669 |
| **CDCA** | 21 | 12.689<br>[2.338 - 18.243] | 23 | 10.69<br>[3.673 - 22.813] | 16 | 0.863<br>[0.342 - 2.475] | 11 | 0.656<br>[0.021 - 1.873] | $2.2 \times 10^{-9}$ | 0.956 | 0.406 |
| **TCDCA** | 21 | 1.262<br>[0.925 - 2.091] | 23 | 0.966<br>[0.518 - 1.596] | 16 | 0.421<br>[0.243 - 0.807] | 11 | 0.33<br>[0.239 - 0.658] | $1.0 \times 10^{-6}$ | 0.142 | 0.597 |
| **UDCA** | 21 | 0.148<br>[0.047 - 0.396] | 23 | 0.166<br>[0.043 - 0.575] | 16 | 0.022<br>[0.009 - 0.058] | 11 | 0.054<br>[0.026 - 0.089] | $3.9 \times 10^{-5}$ | 0.174 | 0.289 |
| **GUDCA** | 15 | 0.005<br>[0.002 - 0.011] | 18 | 0.005<br>[0.003 - 0.0125] | 7 | 0.002<br>[0.001 - 0.004] | 10 | 0.002<br>[0.001 - 0.002] | $2.4 \times 10^{-4}$ | 0.445 | 0.445 |
| **LCA** | 19 | 0.074<br>[0.038 - 0.099] | 23 | 0.089<br>[0.045 - 0.188] | 13 | 0.013<br>[0.007 - 0.0335] | 8 | 0.008<br>[0.002 - 0.029] | $2.8 \times 10^{-7}$ | 0.907 | 0.437 |
| **αMUA** | 20 | 3.759<br>[1.874 - 8.254] | 22 | 4.66<br>[2.045 - 8.023] | 13 | 0.832<br>[0.305 - 1.178] | 7 | 0.948<br>[0.551 - 1.198] | $3.0 \times 10^{-6}$ | 0.524 | 0.978 |
| **TαMUA** | 21 | 1.777<br>[1.060 - 2.147] | 23 | 1.085<br>[0.756 - 1.877] | 16 | 0.706<br>[0.428 - 1.121] | 11 | 0.693<br>[0.338 - 0.882] | $1.0 \times 10^{-5}$ | 0.135 | 0.837 |
| **βMUA** | 18 | 0.834<br>[0.392 - 1.561] | 22 | 0.59<br>[0.196 - 1.794] | 13 | 0.085<br>[0.045 - 0.209] | 7 | 0.134<br>[0.077 - 0.242] | $2.9 \times 10^{-7}$ | 0.858 | 0.150 |
| **THDCA** | 18 | 0.039<br>[0.016 - 0.130] | 18 | 0.102<br>[0.036 - 0.237] | 16 | 0.218<br>[0.144 - 0.376] | 11 | 0.324<br>[0.193 - 0.792] | $2.0 \times 10^{-6}$ | 0.079 | 0.932 |

In control animals on standard diet, exercise tended to increase total bile acids by 4% with the greatest effect observed for unconjugated species (+9%). At the same time, other bile acids decreased, for example conjugated bile acids (−28%) and 12-α-OH (−25%) species. In exercising HFD animals, opposite trends were seen with reductions in total bile acids (−12%), unconjugated bile acids (−40%), primary bile acids (−21%) and 12-α-OH species (−29%). However, the alterations in the bile acid profile of exercising control and HFD animals did not reach statistical significance. Furthermore, no interaction between diet and exercise could be found (Table 1 and Figure 3).

### 3.4. Long term HFD dysregulates bile acid levels in stool that cannot be reverted by exercise

Measurement of primary and secondary bile acids (Suppl. Table S1) in stool showed 2-fold higher total bile acid levels and an alteration of the composition of the faecal bile acid pool in HFD animals when compared to controls (Figure 4). The doubling of total bile acids in stool was the result of similar changes in conjugated and unconjugated species, as well as secondary bile acids in these animals. A particularly strong effect of HFD was seen for 12-α-OH bile acids where faecal levels increased 7.7-fold (Table 2). A detailed analysis of individual bile acids showed that levels of conjugated (both taurine and glycine) cholic acid were increased between 3.8- and 2.8-fold in the HFD group compared to inactive controls. The greatest change was detected for deoxycholic acid (+8.5-fold) and its taurine (+5.4-fold) and glycine (+2.8-fold) conjugated species. Table 2 provides detailed information on faecal bile acid levels in all groups. Similar to the results in serum, exercise did not alter faecal bile acid levels, independently of the diet (Table 2 and Figure 4).

**Figure 4.**
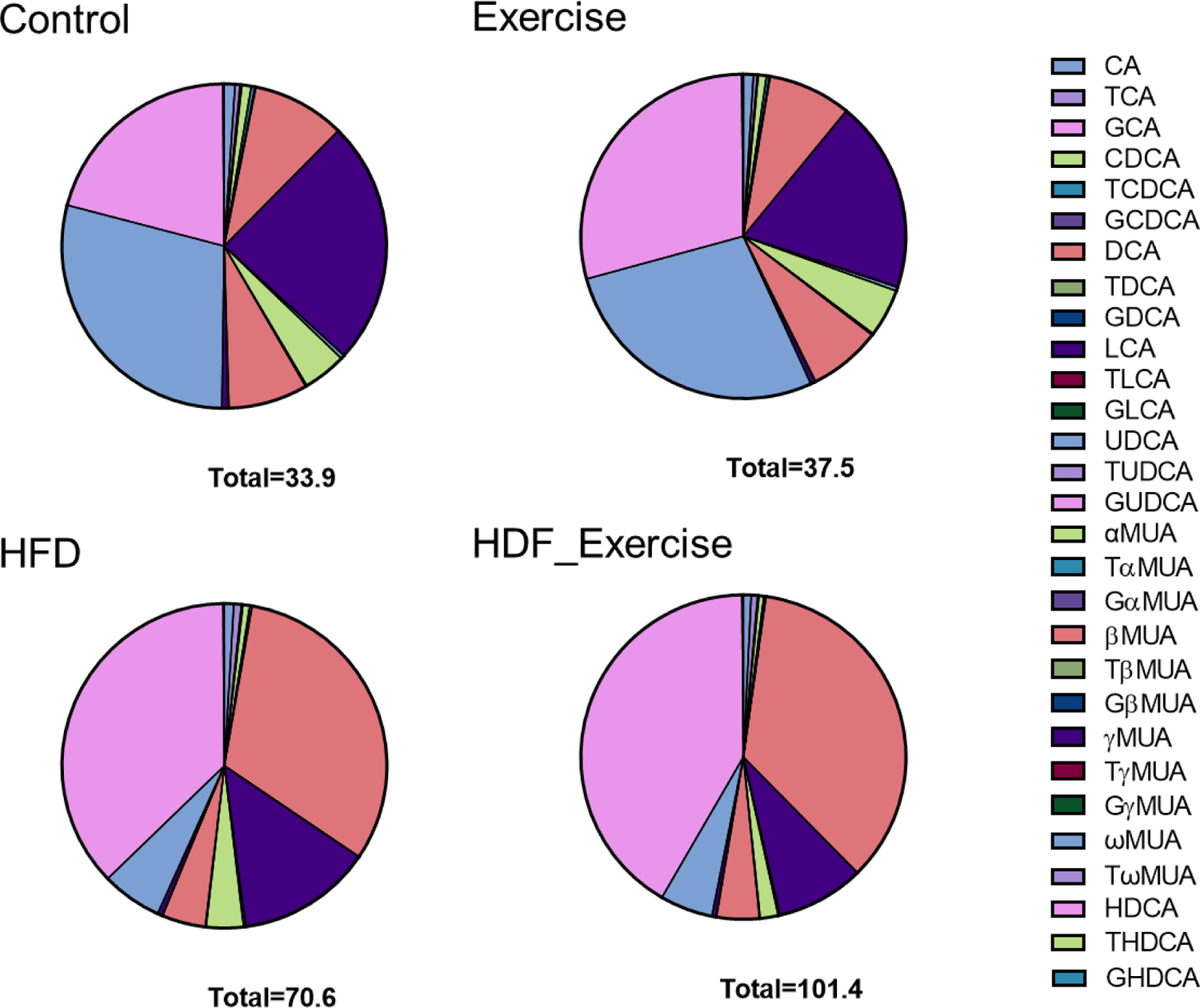
Distribution of bile acids (median) in stool of each of the intervention groups and controls. Total concentration of all species of bile acids is shown at the bottom of each graph.

**Table 2.**
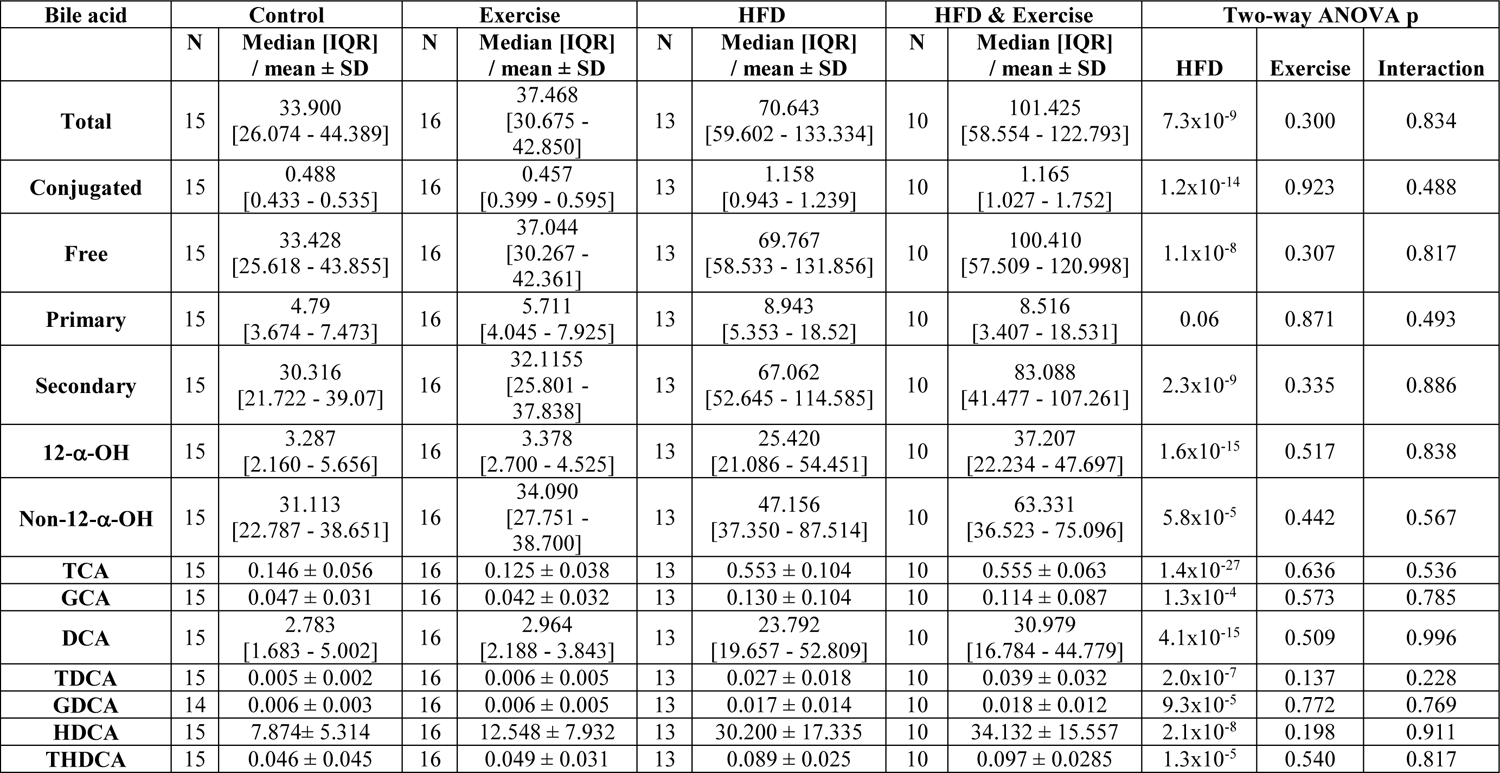
Significant changes in bile acids (µM) in stool associated with high fat diet and physical activity. Normally distributed data is shown as mean ± SD and not normally distributed data is shown as median [interquartile range]. Threshold for significance p =7.0×10^-4^.

| Bile acid | Control |  | Exercise |  | HFD |  | HFD & Exercise |  | Two-way ANOVA p |  |  |
| --- | --- | --- | --- | --- | --- | --- | --- | --- | --- | --- | --- |
| | N | Median [IQR] / mean ± SD | N | Median [IQR] / mean ± SD | N | Median [IQR] / mean ± SD | N | Median [IQR] / mean ± SD | HFD | Exercise | Interaction |
| **Total** | 15 | 33.900<br>[26.074 - 44.389] | 16 | 37.468<br>[30.675 - 42.850] | 13 | 70.643<br>[59.602 - 133.334] | 10 | 101.425<br>[58.554 - 122.793] | $7.3 \times 10^{-9}$ | 0.300 | 0.834 |
| **Conjugated** | 15 | 0.488<br>[0.433 - 0.535] | 16 | 0.457<br>[0.399 - 0.595] | 13 | 1.158<br>[0.943 - 1.239] | 10 | 1.165<br>[1.027 - 1.752] | $1.2 \times 10^{-14}$ | 0.923 | 0.488 |
| **Free** | 15 | 33.428<br>[25.618 - 43.855] | 16 | 37.044<br>[30.267 - 42.361] | 13 | 69.767<br>[58.533 - 131.856] | 10 | 100.410<br>[57.509 - 120.998] | $1.1 \times 10^{-8}$ | 0.307 | 0.817 |
| **Primary** | 15 | 4.79<br>[3.674 - 7.473] | 16 | 5.711<br>[4.045 - 7.925] | 13 | 8.943<br>[5.353 - 18.52] | 10 | 8.516<br>[3.407 - 18.531] | 0.06 | 0.871 | 0.493 |
| **Secondary** | 15 | 30.316<br>[21.722 - 39.07] | 16 | 32.1155<br>[25.801 - 37.838] | 13 | 67.062<br>[52.645 - 114.585] | 10 | 83.088<br>[41.477 - 107.261] | $2.3 \times 10^{-9}$ | 0.335 | 0.886 |
| **12-α-OH** | 15 | 3.287<br>[2.160 - 5.656] | 16 | 3.378<br>[2.700 - 4.525] | 13 | 25.420<br>[21.086 - 54.451] | 10 | 37.207<br>[22.234 - 47.697] | $1.6 \times 10^{-15}$ | 0.517 | 0.838 |
| **Non-12-α-OH** | 15 | 31.113<br>[22.787 - 38.651] | 16 | 34.090<br>[27.751 - 38.700] | 13 | 47.156<br>[37.350 - 87.514] | 10 | 63.331<br>[36.523 - 75.096] | $5.8 \times 10^{-5}$ | 0.442 | 0.567 |
| **TCA** | 15 | $0.146 \pm 0.056$ | 16 | $0.125 \pm 0.038$ | 13 | $0.553 \pm 0.104$ | 10 | $0.555 \pm 0.063$ | $1.4 \times 10^{-27}$ | 0.636 | 0.536 |
| **GCA** | 15 | $0.047 \pm 0.031$ | 16 | $0.042 \pm 0.032$ | 13 | $0.130 \pm 0.104$ | 10 | $0.114 \pm 0.087$ | $1.3 \times 10^{-4}$ | 0.573 | 0.785 |
| **DCA** | 15 | 2.783<br>[1.683 - 5.002] | 16 | 2.964<br>[2.188 - 3.843] | 13 | 23.792<br>[19.657 - 52.809] | 10 | 30.979<br>[16.784 - 44.779] | $4.1 \times 10^{-15}$ | 0.509 | 0.996 |
| **TDCA** | 15 | $0.005 \pm 0.002$ | 16 | $0.006 \pm 0.005$ | 13 | $0.027 \pm 0.018$ | 10 | $0.039 \pm 0.032$ | $2.0 \times 10^{-7}$ | 0.137 | 0.228 |
| **GDCA** | 14 | $0.006 \pm 0.003$ | 16 | $0.006 \pm 0.005$ | 13 | $0.017 \pm 0.014$ | 10 | $0.018 \pm 0.012$ | $9.3 \times 10^{-5}$ | 0.772 | 0.769 |
| **HDCA** | 15 | $7.874 \pm 5.314$ | 16 | $12.548 \pm 7.932$ | 13 | $30.200 \pm 17.335$ | 10 | $34.132 \pm 15.557$ | $2.1 \times 10^{-8}$ | 0.198 | 0.911 |
| **THDCA** | 15 | $0.046 \pm 0.045$ | 16 | $0.049 \pm 0.031$ | 13 | $0.089 \pm 0.025$ | 10 | $0.097 \pm 0.0285$ | $1.3 \times 10^{-5}$ | 0.540 | 0.817 |

### 3.5. Long term HFD affects the bile acid ratios that cannot be reverted by exercise

To identify potential effects of HFD and exercise on bile acid metabolism, ratios between different bile acid species in serum and stool were calculated and compared between groups (Suppl. Table S2). Moreover, serum/stool ratios were calculated for each bile acid as a proxy for their enterohepatic circulation. These ratios were also compared between groups. HFD feeding increased taurine conjugation, but not glycine conjugation. All ratios with taurine conjugated bile acid species in the numerator were markedly higher in both HFD groups. This effect was seen in serum and stool (Table 3). Also, the CA/CDCA ratio in serum was increased in response to HFD.

**Table 3.**
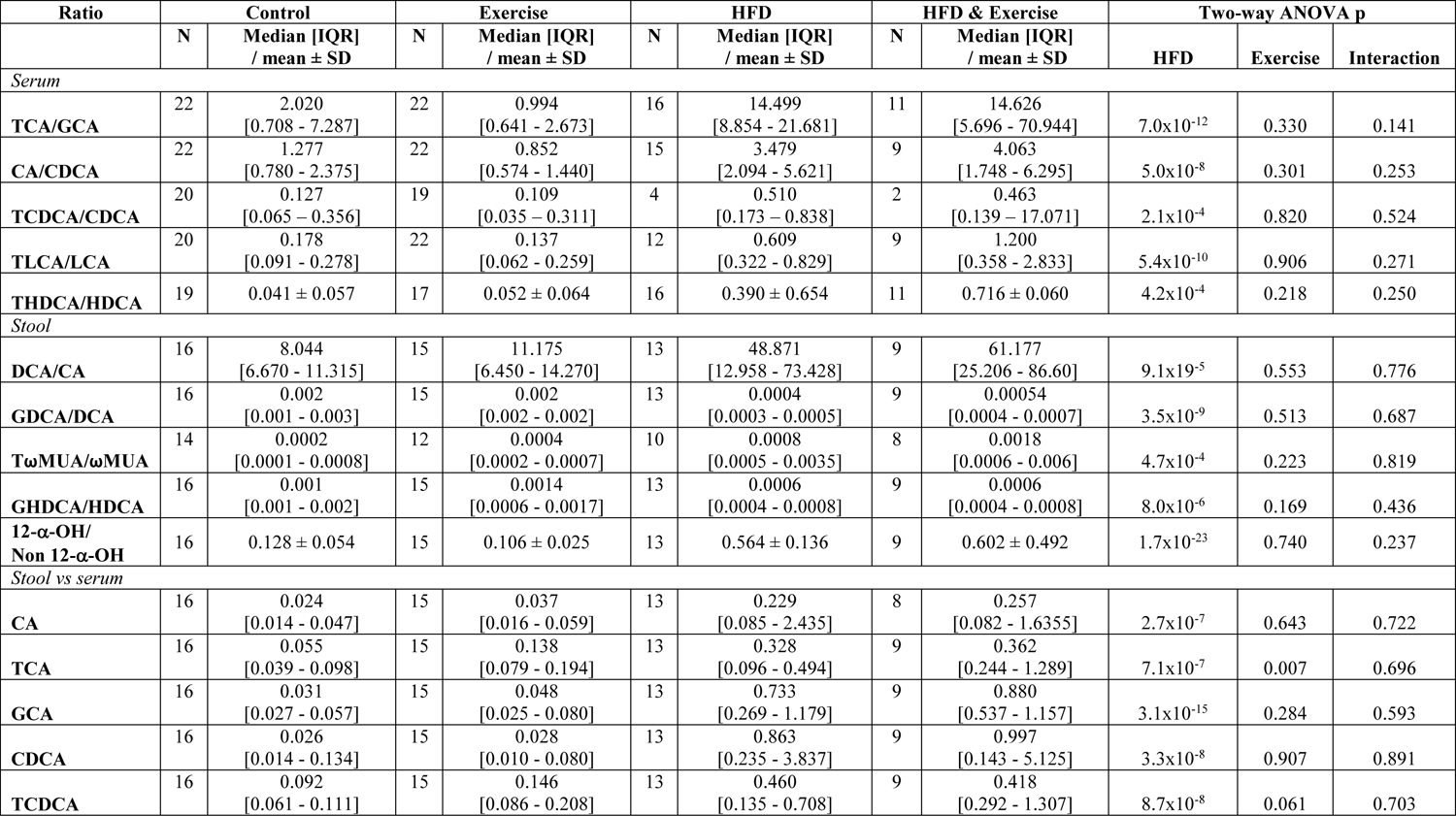
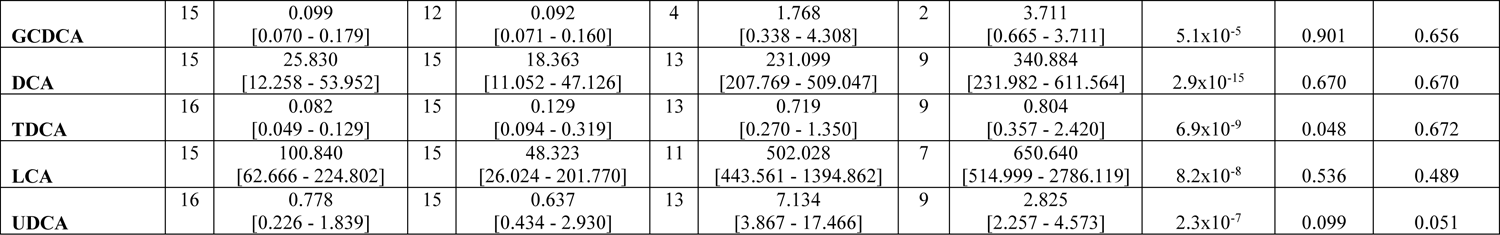
Significant changes in bile acid ratios in serum and stool in response to HDF and exercise (threshold for significance p=6.6×10^-4^)

In stool, the ratio between 12-α-OH and non-12-α-OH bile acids was increased in response to HFD. Furthermore, intestinal deconjugation of CA was markedly increased as shown by a higher faecal DCA/CA ratio. In contrast, glycine conjugation of DCA and HDCA was reduced by HFD in active and sedentary animals compared to control animals on standard diet (Table 3).

Higher stool-to-serum ratios were observed in HFD animals for ten out of the 29 bile acid species when compared to their respective controls on standard diet (Table 3). Moderate exercise did not modify the HFD-related alterations of the bile acid profile in serum and stool and had no impact on enterohepatic circulation of any of the bile acid analyses (Table 3).

### 3.6. Bile acid alterations correlate with bone parameters

Serum bile acids were consistently correlated with structural indices of cortical, but not trabecular bone. The total bile acid concentration in serum was positively related with cortical bone mass (r^2^=0.36). Specifically, GCA, CDCA and a-MCA were consistently correlated with cortical bone parameters (Table 4). In contrast, faecal bile acids were negatively correlated with structural indices of cortical and trabecular bone. The strongest correlations were found for TCA, which correlated inversely with trabecular bone volume, trabecular thickness, cortical thickness, and cortical bone mass with r ranging from −0.656 to −0.805 (Table 4).

**Table 4.** Correlation between bile acids and bile acid ratios in serum and stool and bone parameters obtained by microCT. Values are shown as pearman’s Rho (p value). Significant results are shown in bold (threshold for significance 8.0×10^-5^).

| Bile acid \ uCT parameter | N | BV/TV | BS/TV | TbTh | TbN | CtAr | CtAr /Tt.Ar | CtTh | PsPm | EcPm | Imax | Imin | J |
| --- | --- | --- | --- | --- | --- | --- | --- | --- | --- | --- | --- | --- | --- |
| *Serum levels* |  |  |  |  |  |  |  |  |  |  |  |  |  |
| **Total BAs** | 39 | 0.500<br>(0.001) | 0.484<br>(0.002) | 0.513<br>(0.001) | 0.525<br>(0.001) | 0.556<br>($2.3 \times 10^{-4}$) | **0.602**<br>($5.1 \times 10^{-5}$) | 0.586<br>($8.9 \times 10^{-5}$) | 0.531<br>(0.001) | 0.399<br>(0.012) | 0.384<br>(0.016) | 0.488<br>(0.002) | 0.422<br>(0.007) |
| **Free BAs** | 39 | 0.443<br>(0.005) | 0.426<br>(0.007) | 0.523<br>(0.001) | 0.430<br>(0.006) | 0.582<br>($1.0 \times 10^{-4}$) | 0.564<br>($1.9 \times 10^{-4}$) | **0.609**<br>($3.8 \times 10^{-5}$) | 0.522<br>(0.001) | 0.391<br>(0.014) | 0.342<br>(0.033) | 0.545<br>($3.3 \times 10^{-4}$) | 0.430<br>(0.006) |
| **Primary BAs** | | 0.543<br>($3.5 \times 10^{-4}$) | 0.529<br>($5.4 \times 10^{-4}$) | 0.541<br>($3.7 \times 10^{-4}$) | 0.563<br>($1.9 \times 10^{-4}$) | 0.572<br>($1.4 \times 10^{-4}$) | **0.633**<br>($1.5 \times 10^{-5}$) | **0.609**<br>($3.9 \times 10^{-5}$) | 0.545<br>($3.4 \times 10^{-4}$) | 0.392<br>(0.014) | 0.388<br>(0.0159) | 0.488<br>(0.002) | 0.420<br>(0.008) |
| **Non-12-α-OH BAs** | 39 | 0.456<br>(0.004) | 0.450<br>(0.004) | 0.499<br>(0.001) | 0.469<br>(0.003) | 0.580<br>($1.1 \times 10^{-4}$) | **0.593**<br>($7.1 \times 10^{-5}$) | 0.584<br>($9.5 \times 10^{-5}$) | 0.553<br>($2.6 \times 10^{-4}$) | 0.444<br>(0.005) | 0.419<br>(0.008) | 0.527<br>(0.001) | 0.463<br>(0.003) |
| **GCA** | 38 | 0.539<br>($4.8 \times 10^{-4}$) | 0.550<br>($3.5 \times 10^{-4}$) | 0.586<br>($1.1 \times 10^{-4}$) | 0.553<br>($3.2 \times 10^{-4}$) | **0.619**<br>($3.2 \times 10^{-5}$) | **0.624**<br>($2.8 \times 10^{-5}$) | **0.649**<br>($1.0 \times 10^{-5}$) | **0.641**<br>($1.5 \times 10^{-5}$) | 0.496<br>(0.001) | 0.424<br>(0.008) | **0.599**<br>($7.1 \times 10^{-5}$) | 0.511<br>(0.001) |
| **CDCA** | 38 | 0.480<br>(0.002) | 0.467<br>(0.003) | 0.545<br>($4.0 \times 10^{-4}$) | 0.466<br>(0.003) | **0.631**<br>($2.1 \times 10^{-5}$) | **0.610**<br>($4.9 \times 10^{-5}$) | **0.651**<br>($9.5 \times 10^{-6}$) | 0.575<br>($1.6 \times 10^{-4}$) | 0.437<br>(0.003) | 0.381<br>(0.02) | 0.590<br>($9.7 \times 10^{-5}$) | 0.478<br>(0.002) |
| **αMUA** | 32 | 0.562<br>($8.2 \times 10^{-4}$) | 0.543<br>(0.001) | 0.602<br>($2.6 \times 10^{-4}$) | 0.561<br>($8.3 \times 10^{-4}$) | **0.677**<br>($2.1 \times 10^{-5}$) | **0.708**<br>($5.6 \times 10^{-6}$) | **0.691**<br>($1.2 \times 10^{-5}$) | **0.672**<br>($2.5 \times 10^{-5}$) | 0.516<br>(0.003) | 0.434<br>(0.01) | 0.622<br>($1.4 \times 10^{-4}$) | 0.513<br>(0.003) |
| *Stool levels* |  |  |  |  |  |  |  |  |  |  |  |  |  |
| **Conjugated BAs** | 32 | -0.606<br>($2.3 \times 10^{-4}$) | -0.630<br>($1.1 \times 10^{-4}$) | **-0.680**<br>($1.8 \times 10^{-5}$) | -0.583<br>($4.6 \times 10^{-4}$) | -0.593<br>($3.5 \times 10^{-4}$) | **-0.708**<br>($5.8 \times 10^{-6}$) | **-0.689**<br>($1.3 \times 10^{-5}$) | -0.595<br>($3.3 \times 10^{-4}$) | -0.409<br>(0.020) | -0.291<br>(0.107) | -0.425<br>(0.015) | -0.345<br>(0.053) |
| **12-α-OH BAs** | 32 | -0.579<br>(0.001) | -0.593<br>($3.5 \times 10^{-4}$) | -0.618<br>($1.7 \times 10^{-4}$) | -0.565<br>($7.6 \times 10^{-4}$) | -0.601<br>($2.7 \times 10^{-4}$) | **-0.662**<br>($3.6 \times 10^{-5}$) | **-0.651**<br>($5.5 \times 10^{-5}$) | -0.583<br>($4.6 \times 10^{-4}$) | -0.474<br>(0.006) | -0.384<br>(0.030) | -0.504<br>(0.003) | -0.427<br>(0.015) |
| **TCA** | 32 | -0.618<br>($1.7 \times 10^{-4}$) | **-0.656**<br>($4.6 \times 10^{-5}$) | **-0.750**<br>($7.6 \times 10^{-7}$) | -0.595<br>($3.3 \times 10^{-4}$) | **-0.734**<br>($1.7 \times 10^{-6}$) | **-0.778**<br>($1.6 \times 10^{-7}$) | **-0.805**<br>($2.7 \times 10^{-8}$) | **-0.706**<br>($6.2 \times 10^{-6}$) | -0.541<br>(0.003) | -0.418<br>(0.02) | -0.631<br>($1.1 \times 10^{-4}$) | -0.516<br>(0.002) |
| *Bile acid ratios* |  |  |  |  |  |  |  |  |  |  |  |  |  |
| **TCA/GCA (serum)** | 38 | -0.491<br>(0.002) | -0.500<br>(0.001) | -0.509<br>(0.001) | -0.474<br>(0.003) | **-0.606**<br>($5.5 \times 10^{-5}$) | -0.534<br>($5.5 \times 10^{-4}$) | -0.594<br>($8.5 \times 10^{-5}$) | -0.580<br>($1.4 \times 10^{-4}$) | -0.489<br>(0.002) | -0.439<br>(0.006) | -0.570<br>($1.8 \times 10^{-4}$) | -0.500<br>(0.001) |
| **CA/CDCA (serum)** | 36 | -0.452<br>(0.005) | -0.463<br>(0.004) | -0.515<br>(0.001) | -0.457<br>(0.005) | **-0.686**<br>($3.9 \times 10^{-6}$) | **-0.633**<br>($3.4 \times 10^{-5}$) | **-0.641**<br>($2.6 \times 10^{-5}$) | **-0.665**<br>($9.8 \times 10^{-6}$) | **-0.637**<br>($3.0 \times 10^{-5}$) | -0.543<br>($6.2 \times 10^{-4}$) | **-0.672**<br>($7.3 \times 10^{-6}$) | -0.606<br>($9.0 \times 10^{-5}$) |
| **TLCA/LCA (serum)** | 34 | -0.528<br>(0.001) | -0.520<br>(0.002) | -0.544<br>(0.001) | -0.457<br>(0.007) | **-0.628**<br>($7.0 \times 10^{-5}$) | -0.589<br>($2.5 \times 10^{-4}$) | **-0.647**<br>($3.6 \times 10^{-5}$) | -0.541<br>(0.001) | -0.382<br>(0.026) | -0.416<br>(0.014) | -0.544<br>(0.001) | -0.474<br>(0.005) |
| **TωMUA/ωMUA (stool)** | 26 | -0.683<br>($1.2 \times 10^{-4}$) | **-0.699**<br>($7.1 \times 10^{-5}$) | -0.448<br>(0.02) | -0.679<br>($1.4 \times 10^{-4}$) | -0.411<br>(0.037) | -0.521<br>(0.006) | -0.469<br>(0.016) | -0.449<br>(0.021) | -0.287<br>(0.155) | -0.164<br>(0.424) | -0.289<br>(0.152) | -0.198<br>(0.332) |
| **CA (stool/serum)** | 31 | -0.416<br>(0.020) | -0.473<br>(0.007) | -0.514<br>(0.003) | -0.446<br>(0.012) | -0.521<br>(0.003) | **-0.639**<br>($1.1 \times 10^{-4}$) | -0.617<br>($2.2 \times 10^{-4}$) | -0.517<br>(0.003) | -0.394<br>(0.028) | -0.170<br>(0.359) | -0.430<br>(0.016) | -0.298<br>(0.104) |
| **GCA (stool/serum)** | 31 | -0.604<br>(3.2x10<sup>-4</sup>) | **-0.649**<br>**(7.8x10<sup>-5</sup>)** | -0.615<br>(2.3x10<sup>-4</sup>) | **-0.653**<br>**(6.8x10<sup>-5</sup>)** | -0.615<br>(2.3x10<sup>-4</sup>) | **-0.702**<br>**(1.1x10<sup>-5</sup>)** | **-0.690**<br>**(1.7x10<sup>-5</sup>)** | -0.634<br>(1.3x10<sup>-4</sup>) | -0.443<br>(0.013) | -0.360<br>(0.047) | -0.487<br>(0.005) | -0.407<br>(0.023) |
| **CDCA (stool/serum)** | 31 | -0.469<br>(0.008) | -0.547<br>(0.001) | -0.554<br>(0.001) | -0.511<br>(0.003) | -0.644<br>(9.1x10<sup>-5</sup>) | **-0.699**<br>**(1.2x10<sup>-5</sup>)** | **-0.695**<br>**(1.4x10<sup>-5</sup>)** | **-0.640**<br>**(1.1x10<sup>-4</sup>)** | -0.511<br>(0.003) | -0.312<br>(0.088) | -0.537<br>(0.002) | -0.427<br>(0.017) |
| **DCA (stool/serum)** | 31 | **-0.679**<br>**(2.7x10<sup>-5</sup>)** | **-0.683**<br>**(2.3x10<sup>-5</sup>)** | **-0.673**<br>**(3.3x10<sup>-5</sup>)** | **-0.667**<br>**(4.2x10<sup>-5</sup>)** | **-0.726**<br>**(3.7x10<sup>-6</sup>)** | **-0.771**<br>**(3.9x10<sup>-7</sup>)** | **-0.777**<br>**(2.8x10<sup>-7</sup>)** | **-0.685**<br>**(2.1x10<sup>-5</sup>)** | -0.511<br>(0.003) | -0.471<br>(0.007) | -0.607<br>(2.9x10<sup>-4</sup>) | -0.523<br>(0.003) |
| **LCA (stool/serum)** | 29 | -0.595<br>(0.001) | -0.584<br>(0.001) | -0.544<br>(0.002) | -0.587<br>(0.001) | **-0.674**<br>**(6.1x10<sup>-5</sup>)** | -0.624<br>(3.0x10<sup>-4</sup>) | -0.618<br>(3.6x10<sup>-4</sup>) | -0.644<br>(1.6x10<sup>-4</sup>) | -0.569<br>(0.001) | -0.565<br>(0.001) | -0.658<br>(1.0x10<sup>-4</sup>) | -0.618<br>(3.5x10<sup>-4</sup>) |

Stool/serum ratios of most bile acid species were inversely correlated with structural indices of cortical bone, but only DCA was also significantly associated with indices of trabecular bone. All other bile acid species demonstrated no consisted correlations with trabecular bone (Table 4).

## 4. Discussion

In the present rat model, HFD-related obesity coincided with reduced bone mass and profound alterations of bile acid metabolism. Furthermore, bone quality correlated strongly with the concentration of most bile acids in serum and stool. These correlations were most consistent in cortical bone. Regular moderate exercise, an established approach to preserve metabolism and bone health, did not mitigate the effects of HFD feeding on bone and bile acid metabolism.

As one of the few long-term feeding studies, our results expand existing knowledge on the effects of high nutritional fat intake on bone quality. The rats in the given study were exposed to an excessive nutritional lipid content for a large part of their adult life that corresponds to about 25 years in humans. At the end of the intervention period, the animals showed substantial reductions in cortical and trabecular bone mass, and impaired bone stiffness. The results of previous studies that investigated the role of dietary lipids on bone were inconsistent. While some human studies reported increased bone mineral density (BMD) and reduced fragility fractures in obese women (44, 45), others showed an increased risk of developing osteoporosis (46, 47). Likewise, some preclinical studies in mice demonstrated detrimental effects of HFD on bone structure, mostly at trabecular level (48–51), whereas others observed a positive association between body weight and bone size (52). Also, the impact of HFD on biomechanical properties of bone is not well understood, since opposite results have been reported (48, 52).

HFD feeding of rodents is an established model to mimic obesity in people consuming an excessive amount of animal-derived dietary lipids. Especially when combined with physical inactivity, a high nutritional fat content is associated with increased body weight, visceral fat accumulation and metabolic dysfunction that may promote bone loss or degradation (2). The heterogeneous results of previous animal studies may be explained by a shorter duration and variable composition of the hypercaloric diets used (48–52). In tissues with a slow turnover, such as bone, a rather long exposure to HFD may be required to develop established alterations. The HFD used here aimed to trigger bile acid metabolism, which is essential for the digestion and absorption of fat, but also has broad endocrine effects (reviewed in (15)).

The present results showed marked differences of bile acids in serum and stool between HFD and control animals. Alterations of bile acids in obese individuals (53, 54) and healthy volunteers on a fat-enriched diet (55) are known for some time. With our advanced mass spectrometric method, it was possible to obtain differentiated insights into bile acid metabolism. HFD animals showed a pronounced reduction of unconjugated primary bile acids in serum that was largely driven by CDCA. Variable results were observed for secondary bile acids. UDCA and LCA were reduced in HFD animals, whereas THDCA, a metabolite of MCA and LCA, was about 6-fold higher than in controls. THDCA has been proposed as a regulator of cholesterol (56) and glucose in preclinical models of diabetes (57). In diabetic rats, 12 weeks of HFD feeding also decreased total bile acids, particularly CA, in serum and liver. Non-12-α-hydroxylated species also decreased upon HFD feeding, whereas DCA and 12-α-hydroxylated species did not change (58). Likewise, in the present study serum DCA was not altered by HFD. However, 12-α-hydroxylated bile acids, but not non-12-α-hydroxylated species, were reduced. Such inconsistencies may be due to different animal strains, the presence of diabetes, and the duration of HFD.

In the stool of HFD animals, all groups of bile acid were markedly increased. Previous studies suggest that HFD administration overwhelms the intestinal digestive capacity of fat leading to large amounts of undigested fat and bile acids in the distal digestive tract. Ultimately, these bile acids are lost via the stool and are excluded from enterohepatic circulation (59). Looking at individual bile acid species, the largest increase in faecal excretion was seen for DCA, a cytotoxic bile acid that has been associated with non-alcoholic fatty liver disease, DNA damage, obesity and hepatocellular carcinoma (60). This result is in line with previous studies in healthy humans, patients with fatty liver disease (61, 62) and rats treated with HFD (29, 55, 63, 64). High levels of DCA and its precursor TCA are not only cytotoxic, but may also alter the composition of gut microbiota. In accordance with existing data, faecal excretion of 12-α-hydroxylated bile acids was also increased, which is considered a non-invasive marker of early phase of glucose intolerance that also correlates with cumulative energy intake and visceral adipose tissue (29).

A closer look at the composition of the serum bile acid pool shows a relative abundance of CA over CDCA suggesting a preferential activation of the classic pathway of bile acid synthesis over the alternative pathway. Furthermore, HFD feeding was associated with a preponderant conjugation of primary and secondary bile acids with taurine rather than glycine. This observation, which is supported by a previous study in rats (29), alludes to a preferential use of TCA in lipid absorption and a better intestinal reabsorption than CA. In stool, the 12-α-hydroxylated / non 12-α-hydroxylated ratio, which has been found to be correlated with type II diabetes (61) and liver fat content (65), was 5-fold higher upon HFD treatment. Another striking result was the marked elevation of the DCA/CA ratio in HFD animals, which suggests that dietary fat intake modulates bacterial 7α-dehydroxylation in the gut. Considering the significant bactericidal properties of DCA, HFD could largely reshape the gut microbiome by promoting the conversion of CA into DCA (66).

While bile acids are known as endocrine molecules in glucose and lipid homeostasis, their role in bone metabolism remains largely elusive. Previous studies suggest that they may promote bone formation and prevent bone resorption (21–23). These results are supported by the present study showing significant correlations between bile acids and structural indices of bone health. Correlations were stronger in cortical than in trabecular bone and had opposite directions in serum and stool. The coincidence of reduced bone quality and pronounced alterations of serum and stool bile acids in HFD animals may support further studies to identify any direct or indirect underlying mechanistic link. The observed reduction of unconjugated primary bile acids in serum of HFD animals could disturb the delicate equilibrium between bone formation and bone resorption through an impaired ligand availability for the FXR receptor, which regulates osteoblastic Runx2, Osterix, ERK and β-catenin signalling (21, 67). FXR activation also reduces osteoclastogenesis and inhibits the expression of c-Fos and NFATc1 (21). The lack of FXR signalling also decreases bone mass in respective knockout mice (22). In our study, the preponderance of CA over CDCA was negatively associated with cortical bone structure. This observation supports a potential role of FXR as CDCA is the most effective ligand for this receptor (68–70), far better than CA (70). The correlations observed for αMUA, which does not activate FXR, suggest an involvement of additional pathways, such as VDR signalling (71).

Alterations of bile acids in the stool of HFD animals may also impair bone quality through indirect effects. For example, increased amounts of bile acids in stool with cytotoxic potential may increase permeability of the intestinal mucosa. This facilitates the passage of multiple intestinal toxins and metabolically active compounds across the gut-blood-barrier, so that they can reach the bone via the circulation. Modifications of the gut microbiome (72) and inflammatory processes are additional possibilities (73, 74). For example, TCA has been shown to regulate innate immunity by lowering the expression of cytokines and chemokines in macrophages (73). Furthermore, inflammatory bowel disease is known to promote osteoporosis, probably via TNF-α and other pre-inflammatory cytokines (75).

Regular moderate exercise, a lifestyle factor that is known to preserve bone health, increased trabecular mass, thickness and number, as well as cortical area and endocortical perimeter, in control animals on standard diet. In contrast, exercise did neither alleviate the adverse effects of HFD on bone nor did it alter the bile acid profile in serum or stool. When combined with HFD, exercise reduced femoral stiffness and flexural stress beyond the effect of HFD only.

Previous studies that combined HFD with physical activity showed partial compensatory effects of voluntary wheel running (48, 76), but comparability with the present results is limited due to shorter study duration and substantially different exercise modalities. In contrast to the present study, where exercise frequency, duration and intensity were strictly standardized, these previous studies used voluntary wheel running, which facilitates variations between animals in the same group, but also between studies. For example, obese HFD animals tend to be less active than lean controls resulting in shorter exercise duration and lower intensity. The prolonged exercise and feeding protocol of the present study ensures sufficient time for consolidated adaptive responses in bones, which is not necessarily the case in shorter studies. Unlike previous studies (36, 77), the present results failed to show significant effects of physical activity on bile acid metabolism, regardless of the diet. As the exercise protocol was rather moderate, it cannot be excluded that more intense exercise might have yielded different results. However, the endurance exercise protocol was chosen in order to mimic the activity that people typically perform for fitness and disease prevention. Furthermore, exercise-related changes in bile acids may also be of transient nature and thus were not captured by a single-point measurement at the end of the study.

This study has strengths and limitations that should be considered. The rather long duration of the dietary intervention better mimics obesity in Western societies than shorter treatments used in previous studies (48). Moreover, the exercise protocol was well-controlled and designed on the basis of current guidelines that recommend regular moderate activity to reduce disease risk and mortality. Although it cannot be excluded that variations in exercise type and intensity might have yielded different effects, such protocols would be of limited practical relevance when translated to humans. The comprehensive panel of bile acids that was assessed in serum and stool provides unique insights into the effects of common lifestyle factors on bile acid metabolism. However, measurements have only been performed at the end point so that the dynamics over time were not captured. Future studies should include measurements of bile acids that allow a better distinction between biological variability and treatment effects. Finally, apparent differences between murine and human bile acid metabolism hamper the translation of our results to humans. Therefore, human studies are needed to consolidate the present results.

### 4.1. Conclusion

In conclusion, prolonged consumption of HFD induces quantitative and qualitative alterations of bile acid metabolism that were accompanied by significant bone loss. The strong correlations between bile acids and bone quality may reflect a mechanistic link between bile acid and bone metabolism. Importantly, regular moderate exercise improved trabecular and cortical bone quality in lean controls but did not mitigate the adverse effects of HFD on bone and bile acid metabolism. Future studies should substantiate the potential relationship between bile acid metabolism and bone health and identify the underlying pathways.

## Acknowledgements

We thank Assoc Prof Dr Hubert Scharnagl and Dr Karoline Fechter for the critical discussion of the manuscript.

## References

1. WHO. World Health Organisation. Obesity and overweight [Available from: https://www.who.int/news-room/fact-sheets/detail/obesity-and-overweight.

2. Halade GV, Rahman MM, Williams PJ, Fernandes G. High fat diet-induced animal model of age-associated obesity and osteoporosis. J Nutr Biochem 2010; 21(12):1162–9. doi: 10.1016/j.jnutbio.2009.10.002

3. Cooper AJ, Gupta SR, Moustafa AF, Chao AM. Sex/Gender Differences in Obesity Prevalence, Comorbidities, and Treatment. Curr Obes Rep 2021; 10(4):458–66. doi: 10.1007/s13679-021-00453-x

4. Swinburn BA, Sacks G, Hall KD, et al. The global obesity pandemic: shaped by global drivers and local environments. Lancet 2011; 378(9793):804–14. doi: 10.1016/S0140-6736(11)60813-1

5. Reid IR, Ames R, Evans MC, et al. Determinants of total body and regional bone mineral density in normal postmenopausal women--a key role for fat mass. J Clin Endocrinol Metab 1992; 75(1):45–51. doi: 10.1210/jcem.75.1.1619030

6. Felson DT, Zhang Y, Hannan MT, Anderson JJ. Effects of weight and body mass index on bone mineral density in men and women: the Framingham study. J Bone Miner Res 1993; 8(5):567–73. doi: 10.1002/jbmr.5650080507

7. Ravn P, Cizza G, Bjarnason NH, et al. Low body mass index is an important risk factor for low bone mass and increased bone loss in early postmenopausal women. Early Postmenopausal Intervention Cohort (EPIC) study group. J Bone Miner Res 1999; 14(9):1622–7. doi: 10.1359/jbmr.1999.14.9.1622

8. Compston J. Type 2 diabetes mellitus and bone. J Intern Med 2018; 283(2):140–53. doi: 10.1111/joim.12725

9. Bonds DE, Larson JC, Schwartz AV, et al. Risk of fracture in women with type 2 diabetes: the Women’s Health Initiative Observational Study. J Clin Endocrinol Metab 2006; 91(9):3404–10. doi: 10.1210/jc.2006-0614

10. Starup-Linde J, Frost M, Vestergaard P, Abrahamsen B. Epidemiology of Fractures in Diabetes. Calcif Tissue Int 2017; 100(2):109–21. doi: 10.1007/s00223-016-0175-x

11. Vestergaard P. Discrepancies in bone mineral density and fracture risk in patients with type 1 and type 2 diabetes--a meta-analysis. Osteoporos Int 2007; 18(4):427–44. doi: 10.1007/s00198-006-0253-4

12. Yu EW, Putman MS, Derrico N, Abrishamanian-Garcia G, Finkelstein JS, Bouxsein ML. Defects in cortical microarchitecture among African-American women with type 2 diabetes. Osteoporos Int 2015; 26(2):673–9. doi: 10.1007/s00198-014-2927-7

13. Insull W, Jr. Clinical utility of bile acid sequestrants in the treatment of dyslipidemia: a scientific review. South Med J 2006; 99(3):257–73. doi: 10.1097/01.smj.0000208120.73327.db

14. Lefebvre P, Cariou B, Lien F, Kuipers F, Staels B. Role of bile acids and bile acid receptors in metabolic regulation. Physiol Rev 2009; 89(1):147–91. doi: 10.1152/physrev.00010.2008

15. Houten SM, Watanabe M, Auwerx J. Endocrine functions of bile acids. EMBO J 2006; 25(7):1419–25. doi: 10.1038/sj.emboj.7601049

16. Giannini C, Mastromauro C, Scapaticci S, Gentile C, Chiarelli F. Role of bile acids in overweight and obese children and adolescents. Front Endocrinol (Lausanne) 2022; 13(1011994. doi: 10.3389/fendo.2022.1011994

17. Zhao YX, Song YW, Zhang L, et al. Association between bile acid metabolism and bone mineral density in postmenopausal women. Clinics (Sao Paulo) 2020; 75(e1486. doi: 10.6061/clinics/2020/e1486

18. Liu J, Chen Y, Luo Q. The Association of Serum Total Bile Acids With Bone Mineral Density in Chinese Adults Aged 20-59: A Retrospective Cross-Sectional Study. Front Endocrinol (Lausanne) 2022; 13(817437. doi: 10.3389/fendo.2022.817437

19. Deng D, Pan C, Wu Z, et al. An Integrated Metabolomic Study of Osteoporosis: Discovery and Quantification of Hyocholic Acids as Candidate Markers. Front Pharmacol 2021; 12(725341. doi: 10.3389/fphar.2021.725341

20. Ticho AL, Malhotra P, Dudeja PK, Gill RK, Alrefai WA. Bile Acid Receptors and Gastrointestinal Functions. Liver Res 2019; 3(1):31–9. doi: 10.1016/j.livres.2019.01.001

21. Cho SW, An JH, Park H, et al. Positive regulation of osteogenesis by bile acid through FXR. J Bone Miner Res 2013; 28(10):2109–21. doi: 10.1002/jbmr.1961

22. Zheng T, Kang JH, Sim JS, et al. The farnesoid X receptor negatively regulates osteoclastogenesis in bone remodeling and pathological bone loss. Oncotarget 2017; 8(44):76558–73. doi: 10.18632/oncotarget.20576

23. Li Z, Huang J, Wang F, et al. Dual Targeting of Bile Acid Receptor-1 (TGR5) and Farnesoid X Receptor (FXR) Prevents Estrogen-Dependent Bone Loss in Mice. J Bone Miner Res 2019; 34(4):765–76. doi: 10.1002/jbmr.3652

24. Chiang JY. Bile acids: regulation of synthesis. J Lipid Res 2009; 50(10):1955–66. doi: 10.1194/jlr.R900010-JLR200

25. Monte MJ, Marin JJ, Antelo A, Vazquez-Tato J. Bile acids: chemistry, physiology, and pathophysiology. World J Gastroenterol 2009; 15(7):804–16. doi: 10.3748/wjg.15.804

26. Ridlon JM, Kang DJ, Hylemon PB. Bile salt biotransformations by human intestinal bacteria. J Lipid Res 2006; 47(2):241–59. doi: 10.1194/jlr.R500013-JLR200

27. Dawson PA, Lan T, Rao A. Bile acid transporters. J Lipid Res 2009; 50(12):2340–57. doi: 10.1194/jlr.R900012-JLR200

28. Ocvirk S, O’Keefe SJD. Dietary fat, bile acid metabolism and colorectal cancer. Semin Cancer Biol 2021; 73(347-55. doi: 10.1016/j.semcancer.2020.10.003

29. Yoshitsugu R, Kikuchi K, Iwaya H, et al. Alteration of Bile Acid Metabolism by a High-Fat Diet Is Associated with Plasma Transaminase Activities and Glucose Intolerance in Rats. J Nutr Sci Vitaminol (Tokyo) 2019; 65(1):45–51. doi: 10.3177/jnsv.65.45

30. Ruiz-Gaspa S, Guanabens N, Jurado S, et al. Bilirubin and bile acids in osteocytes and bone tissue. Potential role in the cholestatic-induced osteoporosis. Liver Int 2020; 40(11):2767–75. doi: 10.1111/liv.14630

31. Ruiz-Gaspa S, Dubreuil M, Guanabens N, et al. Ursodeoxycholic acid decreases bilirubin-induced osteoblast apoptosis. Eur J Clin Invest 2014; 44(12):1206–14. doi: 10.1111/eci.12355

32. Chin SH, Kahathuduwa CN, Binks M. Physical activity and obesity: what we know and what we need to know. Obes Rev 2016; 17(12):1226–44. doi: 10.1111/obr.12460

33. Bonaiuti D, Shea B, Iovine R, et al. Exercise for preventing and treating osteoporosis in postmenopausal women. Cochrane Database Syst Rev 2002; 3):CD000333. doi: 10.1002/14651858.CD000333

34. Kohrt WM, Bloomfield SA, Little KD, Nelson ME, Yingling VR, American College of Sports M. American College of Sports Medicine Position Stand: physical activity and bone health. Med Sci Sports Exerc 2004; 36(11):1985–96. doi: 10.1249/01.mss.0000142662.21767.58

35. Pinheiro MB, Oliveira J, Bauman A, Fairhall N, Kwok W, Sherrington C. Evidence on physical activity and osteoporosis prevention for people aged 65+ years: a systematic review to inform the WHO guidelines on physical activity and sedentary behaviour. Int J Behav Nutr Phys Act 2020; 17(1):150. doi: 10.1186/s12966-020-01040-4

36. Danese E, Salvagno GL, Tarperi C, et al. Middle-distance running acutely influences the concentration and composition of serum bile acids: Potential implications for cancer risk? Oncotarget 2017; 8(32):52775–82. doi: 10.18632/oncotarget.17188

37. Morville T, Sahl RE, Trammell SA, et al. Divergent effects of resistance and endurance exercise on plasma bile acids, FGF19, and FGF21 in humans. JCI Insight 2018; 3(15). doi: 10.1172/jci.insight.122737

38. Maurer A, Ward JL, Dean K, et al. Divergence in aerobic capacity impacts bile acid metabolism in young women. J Appl Physiol (1985) 2020; 129(4):768–78. doi: 10.1152/japplphysiol.00577.2020

39. Semeraro MD, Beltrami AP, Kharrat F, et al. The impact of moderate endurance exercise on cardiac telomeres and cardiovascular remodeling in obese rats. Front Cardiovasc Med 2022; 9(1080077. doi: 10.3389/fcvm.2022.1080077

40. Prodinger PM, Burklein D, Foehr P, et al. Improving results in rat fracture models: enhancing the efficacy of biomechanical testing by a modification of the experimental setup. BMC Musculoskelet Disord 2018; 19(1):243. doi: 10.1186/s12891-018-2155-y

41. Singh A, Scholze M, Hammer N. On the influence of surface coating on tissue biomechanics - effects on rat bones under routine conditions with implications for image-based deformation detection. BMC Musculoskelet Disord 2018; 19(1):387. doi: 10.1186/s12891-018-2308-z

42. Amplatz B, Zohrer E, Haas C, et al. Bile acid preparation and comprehensive analysis by high performance liquid chromatography-high-resolution mass spectrometry. Clin Chim Acta 2017; 464(85-92. doi: 10.1016/j.cca.2016.11.014

43. Humbert L, Maubert MA, Wolf C, et al. Bile acid profiling in human biological samples: comparison of extraction procedures and application to normal and cholestatic patients. J Chromatogr B Analyt Technol Biomed Life Sci 2012; 899(135-45. doi: 10.1016/j.jchromb.2012.05.015

44. Khosla S, Atkinson EJ, Riggs BL, Melton LJ, 3rd. Relationship between body composition and bone mass in women. J Bone Miner Res 1996; 11(6):857–63. doi: 10.1002/jbmr.5650110618

45. Tang X, Liu G, Kang J, et al. Obesity and risk of hip fracture in adults: a meta-analysis of prospective cohort studies. PLoS One 2013; 8(4):e55077. doi: 10.1371/journal.pone.0055077

46. Nielson CM, Srikanth P, Orwoll ES. Obesity and fracture in men and women: an epidemiologic perspective. J Bone Miner Res 2012; 27(1):1–10. doi: 10.1002/jbmr.1486

47. Compston JE, Watts NB, Chapurlat R, et al. Obesity is not protective against fracture in postmenopausal women: GLOW. Am J Med 2011; 124(11):1043–50. doi: 10.1016/j.amjmed.2011.06.013

48. Picke AK, Sylow L, Moller LLV, et al. Differential effects of high-fat diet and exercise training on bone and energy metabolism. Bone 2018; 116(120-34. doi: 10.1016/j.bone.2018.07.015

49. Scheller EL, Khoury B, Moller KL, et al. Changes in Skeletal Integrity and Marrow Adiposity during High-Fat Diet and after Weight Loss. Front Endocrinol (Lausanne) 2016; 7(102. doi: 10.3389/fendo.2016.00102

50. Zernicke RF, Salem GJ, Barnard RJ, Schramm E. Long-term, high-fat-sucrose diet alters rat femoral neck and vertebral morphology, bone mineral content, and mechanical properties. Bone 1995; 16(1):25–31. doi: 10.1016/s8756-3282(00)80007-1

51. Cao JJ, Gregoire BR, Gao H. High-fat diet decreases cancellous bone mass but has no effect on cortical bone mass in the tibia in mice. Bone 2009; 44(6):1097–104. doi: 10.1016/j.bone.2009.02.017

52. Silva MJ, Eekhoff JD, Patel T, et al. Effects of High-Fat Diet and Body Mass on Bone Morphology and Mechanical Properties in 1100 Advanced Intercross Mice. J Bone Miner Res 2019; 34(4):711–25. doi: 10.1002/jbmr.3648

53. Gadaleta RM, Garcia-Irigoyen O, Moschetta A. Bile acids and colon cancer: Is FXR the solution of the conundrum? Mol Aspects Med 2017; 56(66-74. doi: 10.1016/j.mam.2017.04.002

54. Haeusler RA, Camastra S, Nannipieri M, et al. Increased Bile Acid Synthesis and Impaired Bile Acid Transport in Human Obesity. J Clin Endocrinol Metab 2016; 101(5):1935–44. doi: 10.1210/jc.2015-2583

55. Wan Y, Yuan J, Li J, et al. Unconjugated and secondary bile acid profiles in response to higher-fat, lower-carbohydrate diet and associated with related gut microbiota: A 6-month randomized controlled-feeding trial. Clin Nutr 2020; 39(2):395–404. doi: 10.1016/j.clnu.2019.02.037

56. Eyssen HJ, De Pauw G, Van Eldere J. Formation of hyodeoxycholic acid from muricholic acid and hyocholic acid by an unidentified gram-positive rod termed HDCA-1 isolated from rat intestinal microflora. Appl Environ Microbiol 1999; 65(7):3158–63. doi: 10.1128/AEM.65.7.3158-3163.1999

57. Zheng X, Chen T, Jiang R, et al. Hyocholic acid species improve glucose homeostasis through a distinct TGR5 and FXR signaling mechanism. Cell Metab 2021; 33(4):791–803 e7. doi: 10.1016/j.cmet.2020.11.017

58. Nakade Y, Kitano R, Sakamoto K, et al. Characteristics of bile acid composition in high fat diet-induced nonalcoholic fatty liver disease in obese diabetic rats. PLoS One 2021; 16(2):e0247303. doi: 10.1371/journal.pone.0247303

59. Wang R, Fan X, Lu Y, Chen D, Zhao Y, Qi K. Dietary acetic acid suppress high-fat diet-induced obesity in mice by altering taurine conjugated bile acids metabolism. Curr Res Food Sci 2022; 5(1976-84. doi: 10.1016/j.crfs.2022.10.021

60. Yoshimoto S, Loo TM, Atarashi K, et al. Obesity-induced gut microbial metabolite promotes liver cancer through senescence secretome. Nature 2013; 499(7456):97-101. doi: 10.1038/nature12347

61. Haeusler RA, Astiarraga B, Camastra S, Accili D, Ferrannini E. Human insulin resistance is associated with increased plasma levels of 12alpha-hydroxylated bile acids. Diabetes 2013; 62(12):4184–91. doi: 10.2337/db13-0639

62. Jiao N, Baker SS, Chapa-Rodriguez A, et al. Suppressed hepatic bile acid signalling despite elevated production of primary and secondary bile acids in NAFLD. Gut 2018; 67(10):1881–91. doi: 10.1136/gutjnl-2017-314307

63. Reddy BS, Mangat S, Sheinfil A, Weisburger JH, Wynder EL. Effect of type and amount of dietary fat and 1,2-dimethylhydrazine on biliary bile acids, fecal bile acids, and neutral sterols in rats. Cancer Res 1977; 37(7 Pt 1):2132–7. doi:

64. Hori S, Abe T, Lee DG, et al. Association between 12alpha-hydroxylated bile acids and hepatic steatosis in rats fed a high-fat diet. J Nutr Biochem 2020; 83(108412. doi: 10.1016/j.jnutbio.2020.108412

65. Iwasaki W, Yoshida R, Liu H, et al. The ratio of 12alpha to non-12-hydroxylated bile acids reflects hepatic triacylglycerol accumulation in high-fat diet-fed C57BL/6J mice. Sci Rep 2022; 12(1):16707. doi: 10.1038/s41598-022-20838-9

66. Kurdi P, Kawanishi K, Mizutani K, Yokota A. Mechanism of growth inhibition by free bile acids in lactobacilli and bifidobacteria. J Bacteriol 2006; 188(5):1979–86. doi: 10.1128/JB.188.5.1979-1986.2006

67. Jeong BC, Lee YS, Bae IH, et al. The orphan nuclear receptor SHP is a positive regulator of osteoblastic bone formation. J Bone Miner Res 2010; 25(2):262–74. doi: 10.1359/jbmr.090718

68. Tu H, Okamoto AY, Shan B. FXR, a bile acid receptor and biological sensor. Trends Cardiovasc Med 2000; 10(1):30–5. doi: 10.1016/s1050-1738(00)00043-8

69. Liu J, Lu H, Lu YF, et al. Potency of individual bile acids to regulate bile acid synthesis and transport genes in primary human hepatocyte cultures. Toxicol Sci 2014; 141(2):538–46. doi: 10.1093/toxsci/kfu151

70. Modica S, Gadaleta RM, Moschetta A. Deciphering the nuclear bile acid receptor FXR paradigm. Nucl Recept Signal 2010; 8(e005. doi: 10.1621/nrs.08005

71. Nishida S, Ishizawa M, Kato S, Makishima M. Vitamin D Receptor Deletion Changes Bile Acid Composition in Mice Orally Administered Chenodeoxycholic Acid. J Nutr Sci Vitaminol (Tokyo) 2020; 66(4):370–4. doi: 10.3177/jnsv.66.370

72. Almer G, Semeraro M, Meinitzer A, et al. Impact of long-term high dietary fat intake and regular exercise on serum TMAO and microbiome composition in female rats. Nutrition and healthy ageing 2023; 1–14. doi: 10.3233/NHA-220198

73. Talebian R, Hashem O, Gruber R. Taurocholic acid lowers the inflammatory response of gingival fibroblasts, epithelial cells, and macrophages. J Oral Sci 2020; 62(3):335–9. doi: 10.2334/josnusd.19-0342

74. Allen K, Jaeschke H, Copple BL. Bile acids induce inflammatory genes in hepatocytes: a novel mechanism of inflammation during obstructive cholestasis. Am J Pathol 2011; 178(1):175–86. doi: 10.1016/j.ajpath.2010.11.026

75. Tilg H, Moschen AR, Kaser A, Pines A, Dotan I. Gut, inflammation and osteoporosis: basic and clinical concepts. Gut 2008; 57(5):684–94. doi: 10.1136/gut.2006.117382

76. McCabe LR, Irwin R, Tekalur A, et al. Exercise prevents high fat diet-induced bone loss, marrow adiposity and dysbiosis in male mice. Bone 2019; 118(20-31. doi: 10.1016/j.bone.2018.03.024

77. Mercer KE, Maurer A, Pack LM, et al. Exercise training and diet-induced weight loss increase markers of hepatic bile acid (BA) synthesis and reduce serum total BA concentrations in obese women. Am J Physiol Endocrinol Metab 2021; 320(5):E864–E73. doi: 10.1152/ajpendo.00644.2020

