## Supplementary material for "Impact of high fat diet and exercise on bone and bile acid metabolism in rats": Suppl Table S1

**Suppl. Table S1.** List of bile acids analysed in serum and stool in the study.

| Bile acid | Type | Species | Modification |
|---|---|---|---|
| Cholic acid (CA) | Primary bile acid | Human and murine | - |
| Taurocholic acid (TCA) | Primary bile acid | Human and murine | Taurine |
| Glycocholic acid (GCA) | Primary bile acid | Human and murine | Glycine |
| Chenodeoxycholic acid (CDCA) | Primary bile acid | Human and murine | - |
| Taurochenodeoxycholic acid (TCDCA) | Primary bile acid | Human and murine | Taurine |
| Glycochenodeoxycholic acid (GCDCA) | Primary bile acid | Human and murine | Glycine |
| Deoxycholic acid (DCA) | Secondary bile acid | Human and murine | - |
| Taurodeoxycholic acid (TDCA) | Secondary bile acid | Human and murine | Taurine |
| Glycodeoxycholic acid (GDCA) | Secondary bile acid | Human and murine | Glycine |
| Ursodeoxycholic acid (UDCA) | Secondary bile acid | Human and murine | - |
| Tauroursodeoxycholic acid (TUDCA) | Secondary bile acid | Human and murine | Taurine |
| Glycoursodeoxycholic acid (GUDCA) | Secondary bile acid | Human and murine | Glycine |
| Lithocholic acid (LCA) | Secondary bile acid | Human and murine | - |
| Taurolithocholic acid (TLCA) | Secondary bile acid | Human and murine | Taurine |
| Glycolithocholic acid (GLCA) | Secondary bile acid | Human and murine | Glycine |
| α-muricholic acid (aMUA) | Primary bile acid | Murine only | - |
| Tauro-α-muricholic acid (TaMUA) | Primary bile acid | Murine only | Taurine |
| Glyco-α-muricholic acid (GaMUA) | Primary bile acid | Murine only | Glycine |
| β-muricholic acid (bMUA) | Primary bile acid | Murine only | - |
| Tauro-β-muricholic acid (TbMUA) | Primary bile acid | Murine only | Taurine |
| Glyco-β-muricholic acid (GbMUA) | Primary bile acid | Murine only | Glycine |
| γ-muricholic acid (gMUA) | Primary bile acid | Murine only | - |
| Tauro-γ-muricholic acid (TgMUA) | Primary bile acid | Murine only | Taurine |
| Glyco-γ-muricholic acid (GgMUA) | Primary bile acid | Murine only | Glycine |
| ω-muricholic acid (oMUA) | Secondary bile acid | Murine only | - |
| Tauro-ω-muricholic acid (ToMUA) | Secondary bile acid | Murine only | Taurine |
| Hyodeoxycholic acid (HDCA) | Secondary bile acid | Human and murine | - |
| Taurohyodeoxycholic acid (THDCA) | Secondary bile acid | Human and murine | Taurine |
| Glycohyodeoxycholic acid (GHDCA) | Secondary bile acid | Human and murine | Glycine |

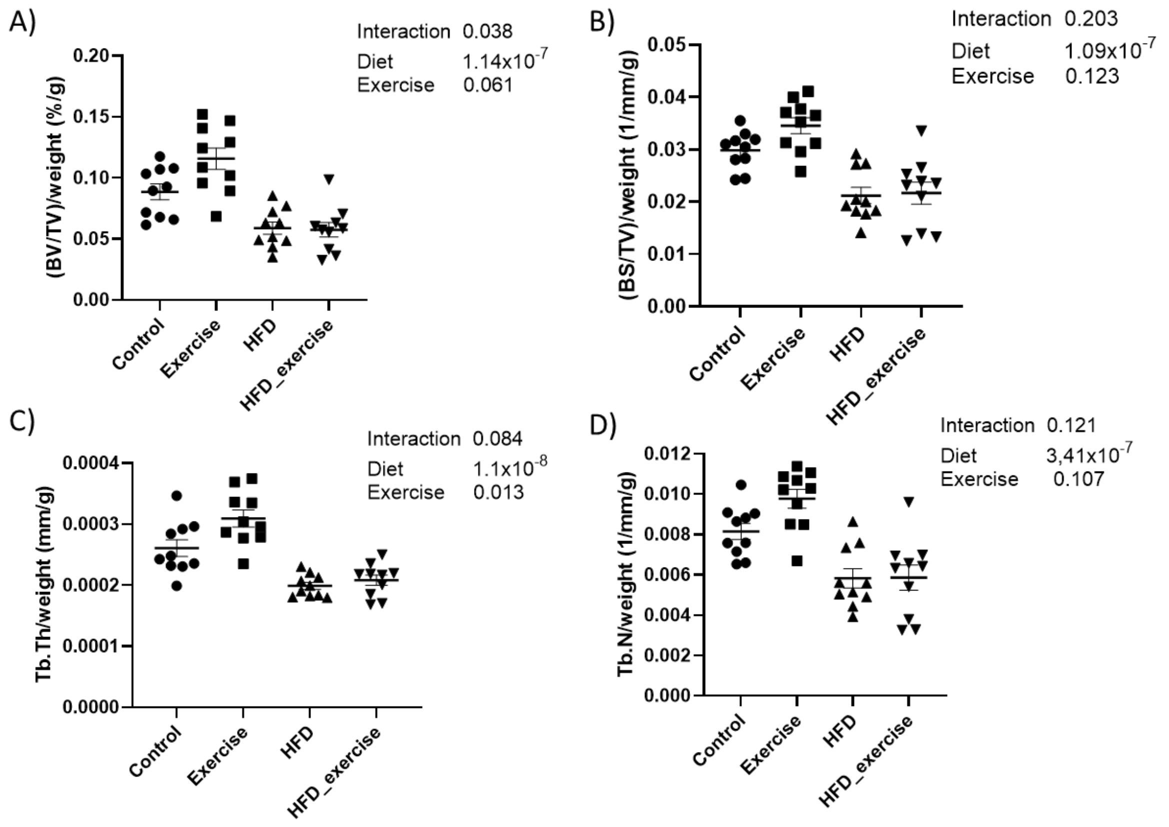

Suppl. Figure S1. Trabecular bone parameters in each group analysed by microCT: A) bone volume fraction (BV/TV); B) bone surface density (BS/TV); C) trabecular thickness (Tb.Th); D) trabecular number (Tb.N). Each data point corresponds to one animal and lines show mean ± SEM. All the parameters are corrected by weight. Two-way ANOVA statistics are shown per graph.

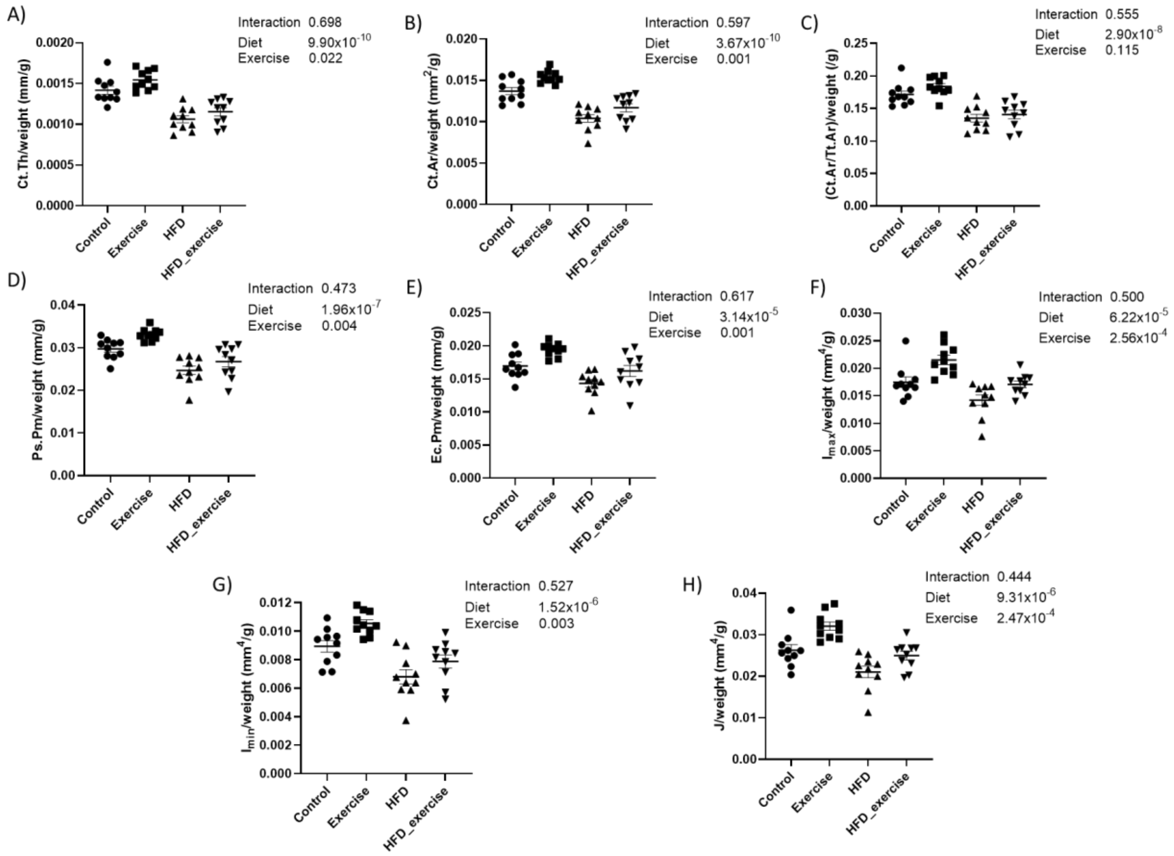

Suppl. Figure S2. Cortical bone parameters in each group analysed by microCT: A) cortical thickness (Ct.Th); B) cortical area (Ct.Ar); C) cortical area fraction (Ct.Ar/Tt.Ar); D) periosteal perimeter (Ps.Pm); E) endocortical perimeter (Ec.Pm); F) maximal inertia ($I_{max}$); G) minimal inertia ($I_{min}$); H) polar moment of inertia (J). Each data point corresponds to one animal and lines show mean ± SEM. All the parameters are corrected by weight. Two-way ANOVA statistics are shown per graph.

**Suppl. Table S2.** List of ratios and bile acid groups analysed in serum and stool.

| **Group** | **Composition** |
|---|---|
| Total BAs | Sum of 29 BAs (Suppl Table S2) |
| Free BAs | CA, CDCA, DCA, UDCA, LCA, αMUA, βMUA, γMUA, ωMUA, HDCA |
| Conjugated BAs | T/G-CA, T/G-CDCA, T/G-DCA, T/G-UDCA, T/G-LCA, T/G-αMUA, T/G-βMUA, T/G-γMUA, TωMUA, T/G-HDCA |
| Primary BAs | As described in Suppl Table S2 |
| Secondary BAs | As described in Suppl Table S2 |
| 12-α-hydroxylated BAs | CA, TCA, GCA, DCA, TDCA, GDCA |
| Non 12-α-hydroxylated | CDCA, TCDCA, GCDCA, UDCA, TUDCA, GUDCA, LCA TLCA, GLCA, αMUA, TαMUA , GαMUA, βMUA, TβMUA, GβMUA, γMUA, TγMUA, GγMUA, ωMUA, TωMUA, HDCA, THDCA, GHDCA |
| **Ratios** | TCA/CA |
| | GCA/CA |
| | TCA / GCA |

| | |
|---|---|
| | CA / CDCA |
| | TCDCA / CDCA |
| | GCDCA / CDCA |
| | TCDCA / GCDCA |
| | DCA / CA |
| | TDCA / DCA |
| | GDCA / DCA |
| | TDCA / GDCA |
| | LCA / CDCA |
| | TLCA / LCA |
| | GLCA / LCA |
| | TLCA / GLCA |
| | UDCA / CDCA |
| | TUDCA / UDCA |
| | GUDCA / UDCA |
| | TUDCA / GUDCA |
| | TαMUA / αMUA |
| | αMUA / CDCA |
| | GαMUA / αMUA |
| | TαMUA / GαMUA |
| | TβMUA / βMUA |
| | GβMUA / βMUA |
| | TβMUA / GβMUA |
| | βMUA / UDCA |
| | TγMUA / γMUA |
| | GγMUA / γMUA |
| | TγMUA / GγMUA |
| | THDCA / HDCA |
| | TωMUA / ωMUA |
| | CA (stool) / CA (serum) |
| | TCA (stool) / TCA (serum) |
| | GCA (stool) / GCA (serum) |
| | CDCA (stool) / CDCA (serum) |
| | TCDCA (stool) / TCDCA (serum) |
| | GCDCA (stool) / GCDCA (serum) |
| | DCA (stool) / DCA (serum) |
| | TDCA (stool) / TDCA (serum) |
| | GDCA (stool) / GDCA (serum) |
| | LCA (stool) / LCA (serum) |
| | TLCA (stool) / TLCA (serum) |
| | GLCA(stool) / GLCA (serum) |
| | UDCA (stool) / UDCA (serum) |
| | TUDCA (stool) / TUDCA (serum) |
| | GUDCA (stool) / GUDCA (serum) |
| | αMUA (stool) / αMUA (serum) |
| | TαMUA (stool) / TαMUA (serum) |
| | GαMUA (stool) / GαMUA (serum) |
| | βMUA (stool) / βMUA (serum) |
| | TβMUA (stool) / TβMUA (serum) |
| | GβMUA (stool) / GβMUA (serum) |
| | γMUA (stool) / γMUA (serum) |
| | ωMUA (stool) / ωMUA (serum) |
| | TωMUA (stool) / TωMUA (serum) |
| | HDCA (stool) / HDCA (serum) |
| | THDCA (stool) / THDCA (serum) |
